# *qad:* An R-package to detect asymmetric and directed dependence in bivariate samples

**DOI:** 10.1101/2022.03.25.485746

**Authors:** Florian Griessenberger, Wolfgang Trutschnig, Robert R. Junker

## Abstract

Correlations belong to the standard repertoire of ecologists for quantifying the strength of dependence between two random variables. Classical dependence measures are usually not capable of detecting non-monotonic or non-functional dependencies. Furthermore, they completely fail to detect asymmetry and direction in dependence, which exist in many situations and should not be ignored. In this paper, we present *qad* (short for quantification of asymmetric dependence), a non-parametric statistical method to quantify directed and asymmetric dependence of bivariate samples. *qad* is applicable in general situations, is sensitive to noise in data, exhibits a good small sample performance, detects asymmetry in dependence, shows high power in testing for independence, requires no assumptions regarding the underlying distribution of the data, and reliably quantifies the information gain/predictability of quantity Y given knowledge of quantity X, and *vice versa* (i.e. *q(X,Y)* ≠ *q(Y, X)*). Here, we briefly recall the methodology underlying *qad*, introduce the functions of the R-package *qad*, which returns estimates for the measures *q*(*X, Y*) denoting the directed dependence of *Y* on *X* (or, equivalently, the influence of *X* on *Y*), *q*(*Y, X*) the directed dependence of *X* on *Y, a*(*X, Y*) ≔ *q*(*X, Y*) − *q*(*Y, X*) the asymmetry in dependence. Furthermore, *qad* can be used to predict *Y* given knowledge of *X*, and *vice versa*. Additionally, we compare empirical performance of *qad* with that of seven other well established measures and demonstrate the applicability of *qad* on ecological datasets. We illustrate that direction and asymmetry in dependence are universal properties of bivariate associations. *qad* thus provides additional information gain and the avoidance of model bias and will therefore advance and facilitate the understanding of ecological systems.

## INTRODUCTION

Although the number of available statistical tools is continuously increasing, classical measures such as correlations often remain the first choice for quantifying the dependence between two random variables (Anderson *et al*. 2021; Bolt *et al*. 2021). Usually, the decision for a specific correlation method is based on the models’ underlying assumptions on the data, e.g., Pearson’s *r* should be used for continuous data, whereas Spearman’s *ρ* is advised for data on the ordinal scale. Both just mentioned dependence measures, however, provide information on different aspects of bivariate distributions: Pearson’s *r* quantifies how linear a relationship is, whereas Spearman’s *ρ* measures the extent of monotonicity. The conclusions drawn from the obtained values have thus take into account the nature of the measures used. Additional insight may be gained by considering other less frequently applied or less well-known dependence measures such as distance correlation (*dCor)* (Székely, Rizzo & Bakirov 2007), a generalization of Pearson correlation and implemented in the R-package *energy*, the information-theoretic-based maximal information coefficient (*MIC)* (Reshef *et al*. 2011) (R-package *minerva*), or the recently developed methods to quantify *asymmetric* dependence *xicor* (Chatterjee 2020) and *qad* (Junker, Griessenberger & Trutschnig 2021).

In recent years, the usefulness of symmetric dependence measures for inferring the structure of complex systems or causality in bi-variate associations has been debated and potential biases have been discussed. For instance, the use of undirected (hence symmetric) dependence measures may lead to inaccurate gene network reconstructions (Wang & Huang 2014) or to misleading conclusions also in equity markets (see, e.g., Okimoto (2008)). Thus, the concept of asymmetry/direction in dependence, which exists in most situations, should not be ignored in data analysis. Whereas in a linear setting the dependence between two variables X and Y is indeed symmetric (Fig. 1A) in the sense that Y can be equally well predicted by knowing X as *vice versa*, the situation, however, is different in general relationships. For instance, for a two-dimensional sample in the form of a parabola (Fig. 1B) or a sinusoidal curve (Fig. 1C), the dependence structure is clearly asymmetric. In these cases, knowing the (value of the) variable *X* strongly improves the predictability of *Y*, whereas in the other direction the information gain is significantly smaller. As an example, consider the year of deglaciation along a glacier forefield and plant diversity. Naturally, the year of deglaciation has a strong influence on plant diversity and a reliable measure of dependence should be able to detect this directed dependence structure. Figure 1D depicts a scatter-plot of the year of deglaciation and plant diversity, established along the forefield of the Ödenwinkelkees glacier (Austria) (see also Junker *et al*. (2020)) together with the results of the directed dependence measure *qad*. Especially, in cases where no a priori knowledge about the causal relationship is available, directional dependence is a useful measure for exploring and estimating the association between two random variables in a more detailed and more realistic way than classical (symmetric/undirected) dependence measures.

**Figure 1:**
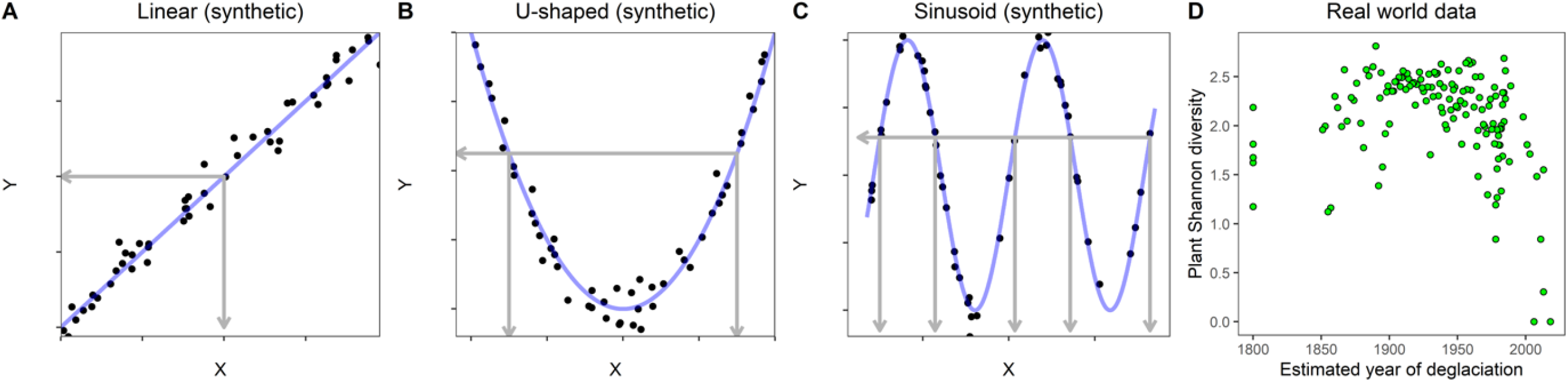
(A-C) Samples of size *n* = 50 drawn from (A) symmetric/undirected as well as (B, C) asymmetric/directed dependence structures. (D) depicts real world data representing plant diversity vs. estimated year of deglaciation at *n* = 140 studied plots. In the symmetric setting (A) the knowledge of *X* provides roughly as much information on *Y* as *vice versa*, whereas in the asymmetric and the real-world data setting (C-D) knowing the value of X allows to predict the value of Y much better than *vice versa*. Asymmetry in dependence is, for instance, detected by the dependence measure *qad*, whereby *Pearson’s r* and *Spearman’s ρ* are not capable of taking into account asymmetry in dependence: **(A)** r(X, Y) = 0.989, ρ(X, Y) = 0.985, qad(X, Y) = qad(Y, X) = 0.861, **(B)** r(X, Y) = 0.051, ρ(X, Y) = 0.021, qad(X, Y) = 0.797 and qad(Y, X) = 0.460, **(C)** r(X, Y) = −0.092, ρ(X, Y) = −0.167, qad(X, Y) = 0.657 and qad(Y, X) = 0.311 and **(D)** r(X, Y) = −0.206, ρ(X, Y) = −0.274 and qad(X, Y) = 0.478 whereas qad(Y, X) = 0.320.

In the era of Big Data network analyses (mostly) based on Pearson’s or more often Spearman’s correlation coefficient are often applied for inferring community structure and (functional) linkages between organisms or individuals, or to understand the microbial, biochemical, and genetic causes for human diseases (Lloyd-Price *et al*. 2019). Despite or maybe because of the fact that undirected and symmetric dependence measures and correlations are standard approaches in network inference, the limitations of these methods have been pointed out just recently (Coenen & Weitz 2018) and directed and asymmetric approaches have been demanded (Amblard & Michel 2011; Karmon & Pilpel 2016; Carr *et al*. 2019).

Here, we present the method *qad*, a non-parametric and directed (hence asymmetric) measure of dependence, which is publicly available in the free software environment R (Griessenberger *et al*. 2021; Junker, Griessenberger & Trutschnig 2021). *qad* returns estimates for the measures *q*(*X, Y*) denoting the directed dependence of *Y* on *X* (or, equivalently, the influence of *X* on *Y*), *q*(*Y, X*) the directed dependence of *X* on *Y*, and *a*(*X, Y*) ≔ *q*(*X, Y*) − *q*(*Y, X*) the asymmetry in dependence. The measure *a*(*X, Y*) for asymmetry in dependence can be interpreted as the difference of the predictability of *Y* given knowledge on *X* and the predictability of *X* given knowledge on *Y*. In this paper, we first describe the methodology of *qad* and demonstrate the application of the R-package *qad*. Furthermore, we compare the empirical performance of *qad* with existing publicly available dependence measures and highlight the information gain by considering asymmetry and direction in dependence. A complementary R-shiny app is available as supporting information facilitating the interpretation and comparison of the results and performance returned by *qad* and other dependence measures. An application of *qad* to real world data concludes the paper. We hope that this introduction to *qad* and the executed comparative analyses as well as the resources provided will be helpful for ecologists and researchers from other disciplines.

## BRIEF METHODOLOGICAL DESCRIPTION OF THE COPULA-BASED DEPENDENCE MEASURE *qad*

Copulas are (bivariate) distribution functions with uniformly distributed univariate marginals. According to Sklar’s theorem (see, for instance, Nelsen (2007)), copulas suffice to capture all scale-invariant dependence between pairs of random variables. The copula-based dependence measure *qad*, originally introduced as *ζ*_1_ in Trutschnig (2011), is defined as a type of distance between the conditional distribution functions of the copula underlying the random vector (*X, Y*) and the uniform distribution representing independence of X and Y. In other words, *qad* measures how much the dependence structure of (*X, Y*) differs from independence. Contrary to many other approaches, qad is able to detect both complete dependence (i.e., Y is a function of X) as well as independence. The method works as follows: Given a two-dimensional sample (*x*_1_, *y*_1_), …, (*x*_*n*_, *y*_*n*_) of size *n* from the random vector (*X, Y*) (see Fig. 2A) the normalized ranks of the sample are calculated first (i.e., we get values of the form (*i*/*n, j*/*n*) for *i, j* ∈ (1, …, *n*)). Then the so-called empirical copula *Ê*_*n*_ is computed (see Fig. 2B). As next step the empirical copula is aggregated to the empirical checkerboard copula (2-dimensional histogram in the copula setting). In fact, the masses of the small squares (empirical copula) are summed up to the larger *N* × *N* squares, whereby the resolution *N* depends on the sample size *n* (see Fig. 2C and D). Note that by default the resolution of the empirical checkerboard copula is proportional to the square root of the sample size; thus, as for any statistical method, *qad* results become more reliable as the sample size increases. We recommend a sample size of no smaller than *n* = 16, resulting in a resolution of *N* = 4. Finally, the conditional distribution functions of the checkerboard copula are compared with the distribution function of the uniform distribution on the unit interval (in the sense that the area between the graphs is calculated). This step is conducted both for the vertical strips (to calculate the influence of *X* on *Y*) as well as the horizontal strips (for the influence of *Y* on *X*), see, for instance, Fig. 2E and F. Computing the sum of all areas and normalizing appropriately with the constant 3 (see Junker, Griessenberger and Trutschnig (2021)) yields the two directed *qad*-values *q*(*X, Y*) ∈ [0,1], quantifying the influence of *X* on *Y* and *q*(*Y, X*) ∈ [0,1], denoting the influence of *Y* on *X*. High values indicate strong associations whereas low values describe weakly dependent random variables. Since low values can also point towards independence, a permutation test is conducted to obtain a *p*-value for *q*(*X, Y*) and *q*(*Y, X*) in testing for independence, i.e., testing the hypothesis *H*_0_: *q*(*X, Y*) = 0 = *q*(*Y, X*).

**Figure 2.**
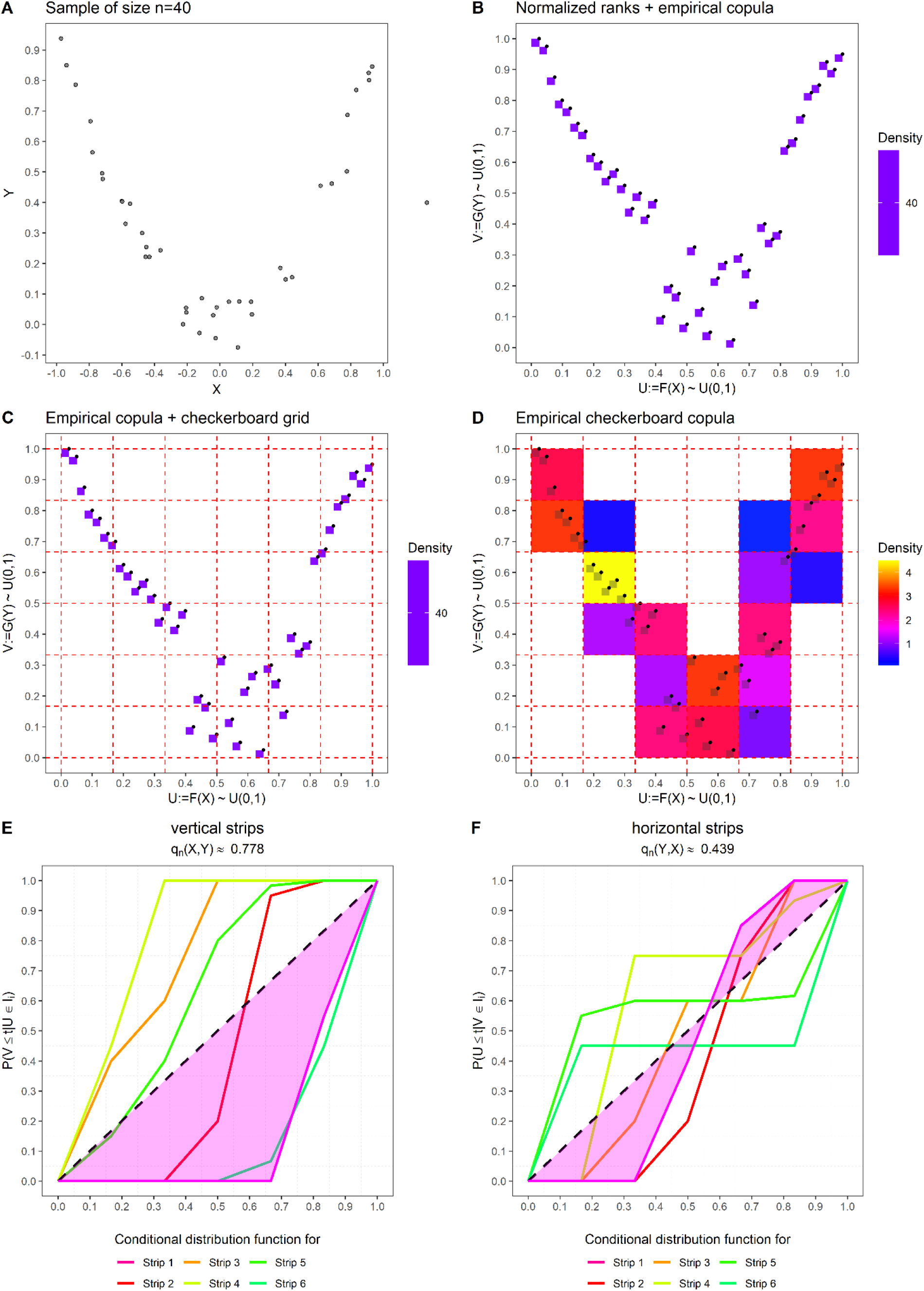
Illustration of the methodology of *qad*. **(A)** Sample of size *n* = 40 drawn from a slightly noisy U-shaped function. **(B)** Empirical copula and normalized ranks (points). Notice that the masses are uniform on each squares and that, by construction of the empirical copula, the upper right corner of the squares are the normalized ranks. **(C)** Empirical copula and the checkerboard grid with resolution *N* = 6. **(D)** Checkerboard aggregation. **(E)** Distance between the conditional distribution functions of the checkerboard copula and the uniform distribution representing independence, for vertical strips (magenta area depicting the distance for one strip) and **(F)** for horizontal strips.

Furthermore, if we have *q*(*X, Y*) > *q*(*Y, X*), then the *qad* estimator informs us that the variable *X* provides more information about *Y* than *vice versa*. The same holds for the reverse direction. This information is also gathered in the measure for asymmetry, which is computed as *a*(*X, Y*) ≔ *q*(*X, Y*) − *q*(*Y, X*) and can therefore attain values within the interval (−1,1). Additionally, as a rank-based quantity *qad* is robust to outliers and invariant with respect to monotone transformations, for instance, log-transformations.

## APPLICATION OF THE R PACKAGE *qad*

The package *qad* is implemented in the software R (R Development Core Team 2020) and is publicly available on CRAN (https://cran.r-project.org/web/packages/qad/index.html). The development version of *qad* is accessible via GitHub (https://github.com/griefl/qad). In the following we briefly sketch the main functions of the package. Additionally, each function contains examples in the description, which are called via the R-help function (e.g., ?qad).

### Calculating the directed dependence measure q

Given bivariate observations (*x*_1_, *y*_1_), …, (*x*_*n*_, *y*_*n*_) of size *n* the function *qad(…)* computes the dependence values *q*(*X, Y*), *q*(*Y, X*), the maximum dependence (i.e. *max(c(q(X,Y), q(Y,X)))*), and the asymmetry in dependence *a*(*X, Y*). The implemented method *qad(…)* requires two numeric vectors containing the observations of the sample, or, alternatively, accepts a numeric data frame of the form *data*.*frame(sample_X, sample_Y)*. The optional argument *p*.*value* (default is TRUE) allows to calculate *p*-values (based on permutations with *nperm* runs) for *q(X,Y)* and *q(Y,X)*. A *p*-value below 0.05 strengthens the hypothesis that *X* and *Y* are not independent. The output of qad shows the dependence values and their respective *p*-values as well as further descriptive statistics, e.g., sample size and the number of unique ranks, which are essential in calculating the resolution of the underlying empirical checkerboard copula.

(Sketch of the output): Data: x1:=x, x2:=y Sample Size: 100 Number of unique ranks: x1: 100; x2: 100; (x1,x2): 100 Resolution: 10 x 10 Dependence measures: ```
|  | *q* | *p.value* |
| --- | --- | --- |
| $q(x_1, x_2)$ | 0.588 | 0.000 |
| $q(x_2, x_1)$ | 0.350 | 0.034 |
| max.dependence | 0.588 | 0.000 |
| asymmetry | 0.237 |  |

Furthermore, the function qad returns an object of class ‘qad’, that allows the application of the generic functions *coef(), summary()* and *plot()*. The plot function generates a two-dimensional histogram (heatmap) visualizing the empirical checkerboard copula. Setting the optional parameter *copula* to FALSE yields a two-dimensional histogram of the unscaled (raw) data.

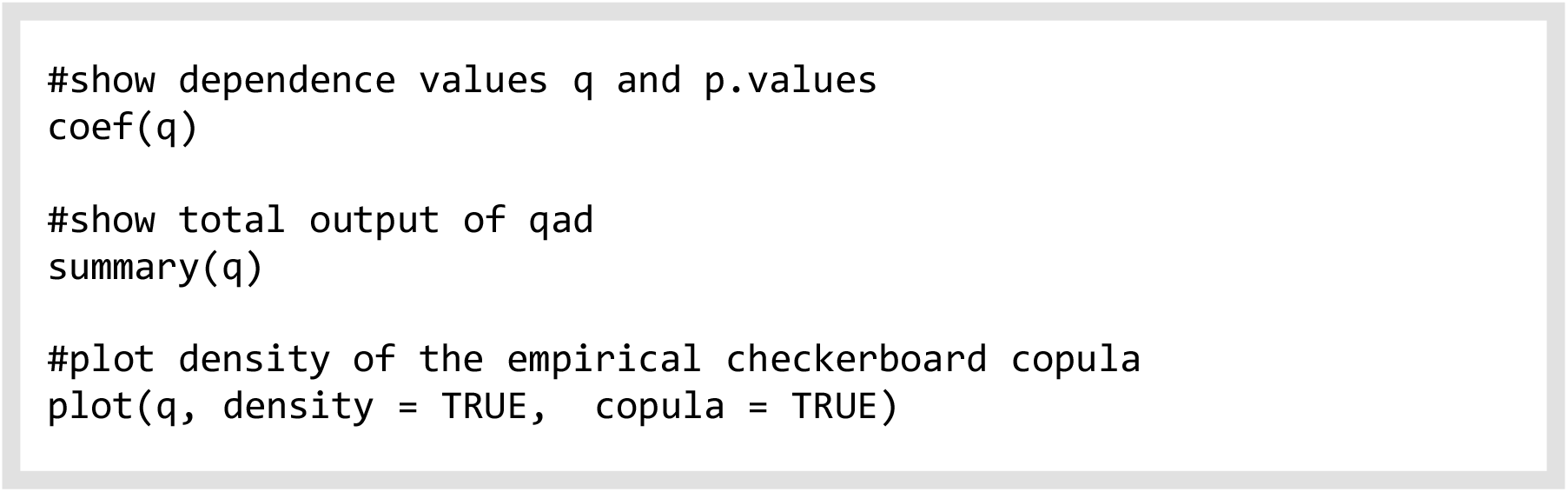

### Using qad as prediction tool

As a by-product of the checkerboard approach, the random variable Y given *X*=*x* and *X* given *Y*=*y* can be predicted for every *x*∈*Range*(*X*) and *y*∈*Range*(*Y*). This additional feature is implemented in the R-function *predict*.*qad(…)*. Note that prediction is possible only within the range of measured X- and Y-values; since *qad* is calculated independently of a parametric regression function, no extrapolation is possible. The function *predict*.*qad(…)* requires three arguments: a ‘qad’ object, the conditioning variable and its values. Then the function returns the probabilities of the event that Y falls into the interval

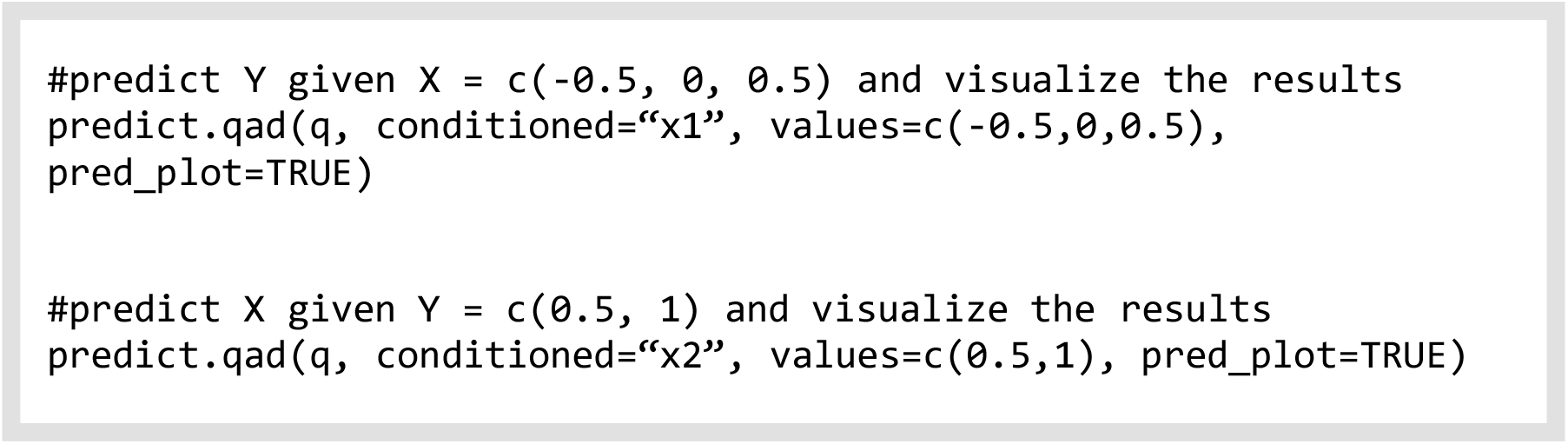

*I*_*j*_ given X=x, or *vice versa*. Via several optional parameters size and numbers of the prediction intervals as well as visualizations may be adjusted as desired.

### Multivariate application of qad

Given a multivariate distribution with more than two variables the function *pairwise*.*qad(…)* can be applied to quantify all pairwise dependencies and allows an interpretation similar to that of a correlation matrix. The method *pairwise*.*qad(…)* requires an *n* × *d*-dimensional numeric matrix, or alternatively, a data.frame of the form *data*.*frame(sample_X1, sample_X2, …*., *sample_Xd)*, describing the observations of a d-dimensional random vector. Among other details, the main output of *pairwise*.*qad(…)* is a data.frame containing all pairwise dependencies and corresponding *p*-values, which may be readily visualized by *heatmap*.*qad()*. Optional parameters allow to select between the dependence or asymmetry values and to highlight all significant pairs.

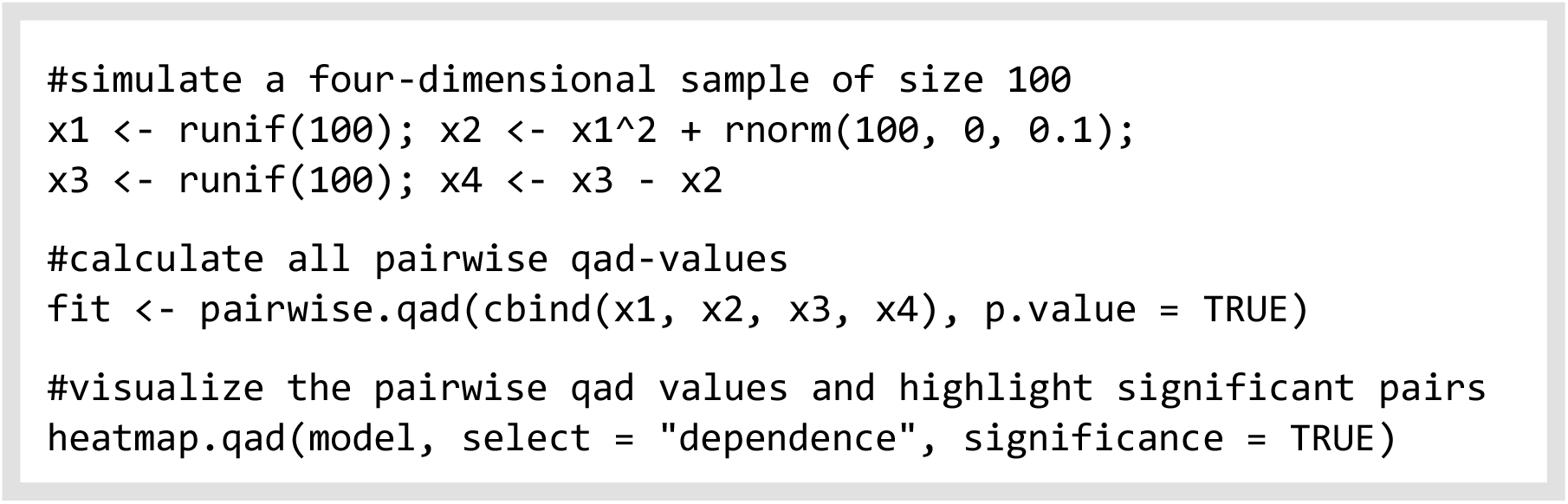

Each of the functions provide several parameters that enables specific adjustments and modifications. For this purpose, we refer to the R-documentation (Griessenberger *et al*. 2021) or the vignette available, e.g., using the following lines of code:

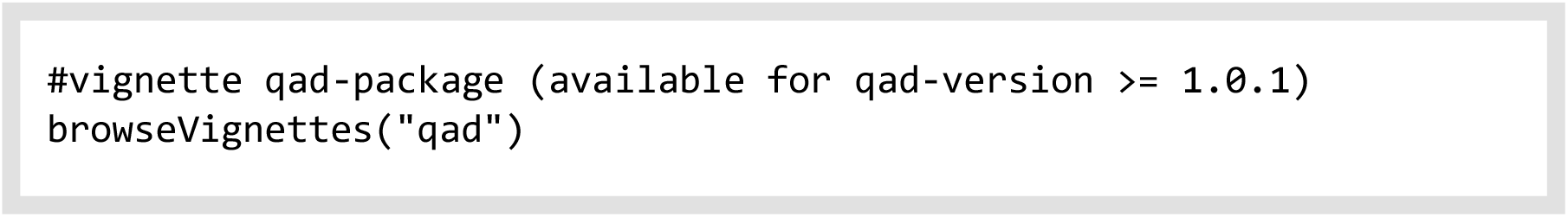

## PERFORMANCE AND COMPARISON OF *qad* WITH OTHER DEPENDENCE MEASURES

Dependence measures such as *qad* that capture the dependence in general situations should assign similar scores of dependence to equally noisy data in a manner independent of the concrete functional relationship (Reshef *et al*. 2011). Accordingly, the dependence *q* between two random variables decreases with increasing noise irrespective of the functional relationship between X and Y (Fig. 3). In a linear setting, *qad* returned dependence values *q* closer to Pearson’s *r*, Spearman’s ρ or distance correlation than other measures of dependence, e.g., the maximal information coefficient *MIC* (Reshef *et al*. 2011), a symmetric dependence measure defined for general situations or *xicor*, a recently introduced new coefficient of correlation (Chatterjee 2020), see Fig. 3A. Unlike commonly used measures of association like Pearson’s *r*, Spearman’s *rho*, or more recent measures such as distance correlation and *MIC*, which are symmetric measures by construction, *qad* (as well as *xicor*) indicated asymmetry in dependence in settings in which (on average) more information on Y could be obtained by knowing the value of X than *vice versa*, i.e., *q*(*X, Y*) > *q*(*Y, X*) (Fig 3B, D, E, F, I). Further empirical studies show that qad ranked high in both runtime analysis and power analysis compared to all other studied dependence measures. Details and further discussions on the results shown in Fig. 3, a runtime evaluation of the different methods, and a power analysis can be found in Supporting information 3. The main features of *qad* compared with the seven other well established dependence measures attaining values in [0,1] or [-1,1], respectively, are also summarized in Table 1. Additionally, to facilitate the applicability and interpretation of the dependence measures, we provide an R-script as well as an R-shiny app allowing the user to evaluate the effects of sample size, noise, and dependence structure on the results obtained by each of the eight dependence measures (see Supplementary information 2: dep_measures.R and app.R).

**Table 1.** Comparison of features of eight well established dependence measures: *qad* (described here and in Junker, Griessenberger and Trutschnig (2021), distance correlation (dCor; Székely, Rizzo and Bakirov (2007)), maximal information coefficient (MIC or MICe; Reshef *et al*. (2011)), robust copula dependence (RCD; Ding *et al*. (2017)), randomized dependence coefficient (rdc; Lopez-Paz, Hennig and Schölkopf (2013)), xicor (Chatterjee 2020) and the commonly used correlation measures by Pearson and Spearman. For each of the eight dependence measures considered here important properties affecting the functionality and the interpretation of the measures are listed.

|  | R-function | Functional types of detection |  |  | Scale invariance | Estimator in [0, 1] | Asymmetry /Directional |
| --- | --- | --- | --- | --- | --- | --- | --- |
|  |  | linear | monotonous | non-monotonous |  |  |  |
| dCor | `energy::dcor()` | ✓ | ✓ | ✓ | ✗ | ✓ | ✗ |
| MICe | `minerva::MIC(est = "mic_e")` | ✓ | ✓ | ✓ | ✓ | ✓ | ✗ |
| Pearson's r | `cor()` | ✓ | ✗ | ✗ | ✗ | ✓* | ✗ |
| qad | `qad::qad(...)` | ✓ | ✓ | ✓ | ✓ | ✓ | ✓ |
| RCD | `rcd::rcd(...)` | ✓ | ✓ | ✓ | ✓ | ✓ | ✗ |
| rdc | 5 lines R-code (see Lopez-Paz *et al.*, 2013) | ✓ | ✓ | ✓ | ✓ | ✓ | ✗ |
| Spearman's $\rho$ | `cor(..., method = "spearman")` | ✓ | ✓ | ✗ | ✓ | ✓* | ✗ |
| xicor | `XICOR::xicor()` | ✓ | ✓ | ✓ | ✓ | ✗ | ✓ |

\* if absolute values are considered which is (in this case) essential to assure comparability with the other measures.

**Figure 3.**
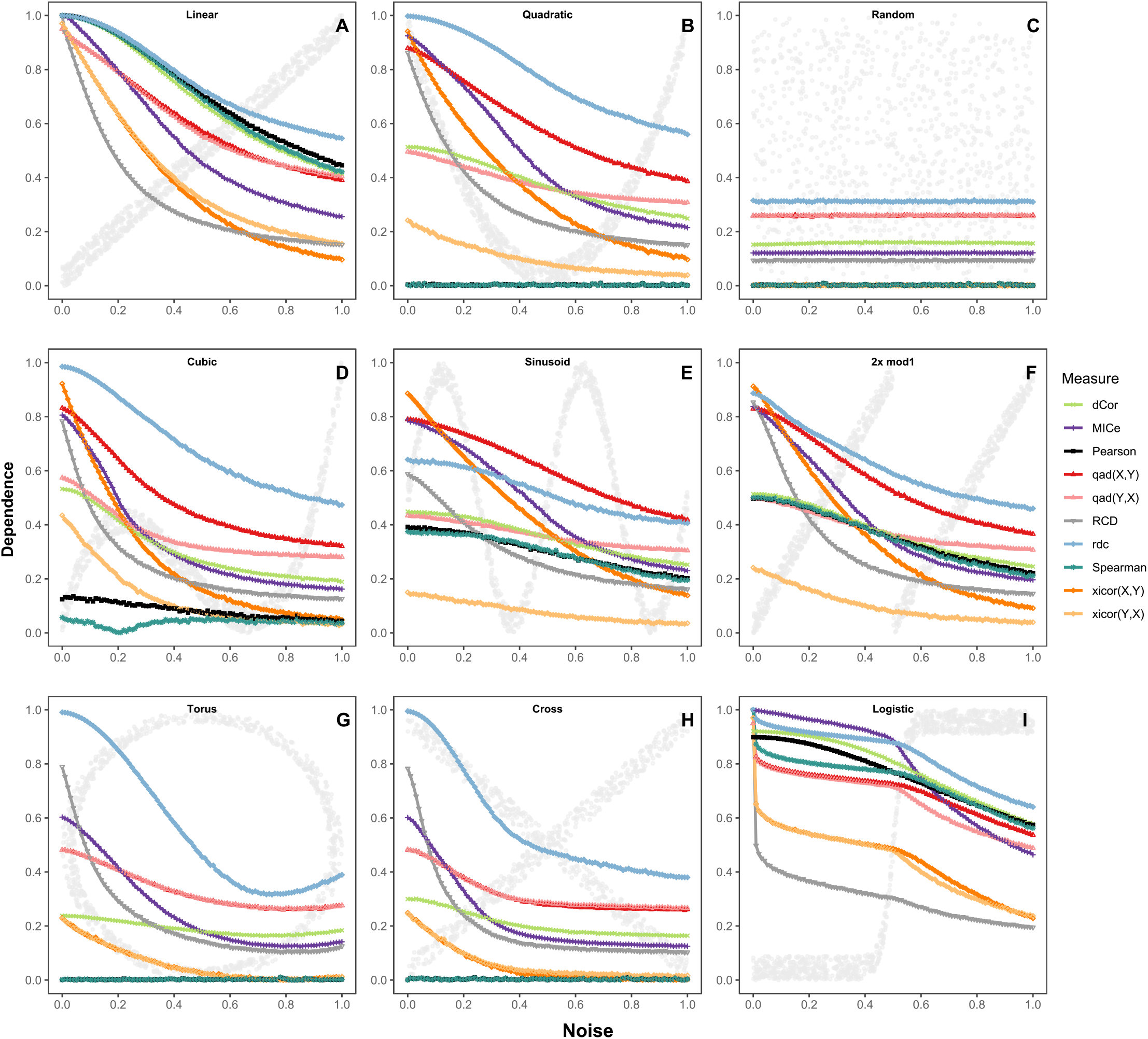
Application of *Pearson’s* |*r*|, *Spearman’s* |*ρ*|, *dCor, MICe, qad, RCD, rdc* and *xicor* to samples of size n = 100 for various kinds of functional and non-functional relationships with vertically added noise (x-axis). **(A-I)** The grey points in the panel backgrounds depict samples of size n = 1000 from the corresponding dependence structures (with noise a = 0.05). Absolute average values for R = 1000 repetitions per noise level are depicted. Note that the dependence structures depicted in (B), (D), (E) and (F) are asymmetric, which is reflected by the two different *qad* and *xicor* values. The other measures of dependence are not able to provide information about asymmetry in dependence.

## APPLYING *qad* ON ECOLOGCIAL DATA

We tested the *qad*-package on a dataset of microbiota and additional environmental metadata publicly available at http://ocean-microbiome.embl.de/companion.html (de Vargas *et al*. 2015; Sunagawa *et al*. 2015; Villar *et al*. 2015; Albanese *et al*. 2018). More precisely, we used the aggregated version of the annotated 16S _mi_tags OTU count table, available in the additional materials of (Albanese *et al*. 2018) and conducted a similar analysis. We computed all pairwise *q*-values across the relative abundances of genera with less than 10% ties and the environmental variables (mean depth, mean salinity, mean temperature, and mean oxygen level), resulting in 94 variables and *n* = 115 samples and compared the results with the values of Pearson’s and Spearman’s correlation coefficient, and illustrated the information gain provided by *qad* over the classical symmetric methods by the number of detected relationships and some specific examples.

Overall, the measure *qad* returned 2907 significant relationships, whereas Spearman’s *ρ* (2564) and Pearson’s *r* (1729) found substantially fewer significant pairs. Furthermore, the classical measures *r* and *ρ* assigned relatively low dependence scores to many relationships that were highly ranked by the measure *qad* (see Fig. 4A, F). This again results from the fact that the classical measures fail to detect many non-linear and non-monotonic dependence structures. Demonstrating major differences in the information gain between *Pearson’s* and *Spearman’s* correlation coefficient in comparison with an asymmetric measure of dependence, we depicted several pairs of variables attaining a high *qad* value but at the same time a low *Pearson* and *Spearman* correlation. For instance, *qad* detected a significant asymmetric dependence between the variable Methylophilaceae-OM43 clade (variable X) and a *Sphingomonas* strain (variable Y), whereas Pearson’s correlation returned a non-significant dependence. The scatterplot depicted in Fig. 4D reveals an inverted U-shaped pattern of the data points, i.e., knowing the relative abundance of Methylophilaceae-OM43 clade is more informative for the prediction of the *Sphingomonas* strain than *vice versa*. Moreover, *qad* also picked up a highly asymmetric dependence structure between Alteromonadaceae-SAR92 clade and a *Marinoscillum* strain. The detected dependence structure can be revealed by a log-transformed scatterplot (Fig. 4E). Note, that *qad* is scale-invariant and hence invariant with respect to log-transformation of samples. We obtained similar results, e.g., for the variables Methylophilaceae-OM43 clade and Alcaligenaceae-MWH-UniP1, see Fig. 4I, and the variables Methylophilaceae-OM43 clade and mean temperature in °C (M_temp), depicted in Fig. 4J. Additionally, Pearson’s *r* reacts very sensitive to outliers (see, for instance, Fig. 4C), which explains that there are several highly ranked relationships found by Pearson’s correlation but ignored by *qad* or Spearman’s correlation.

**Figure 4:**
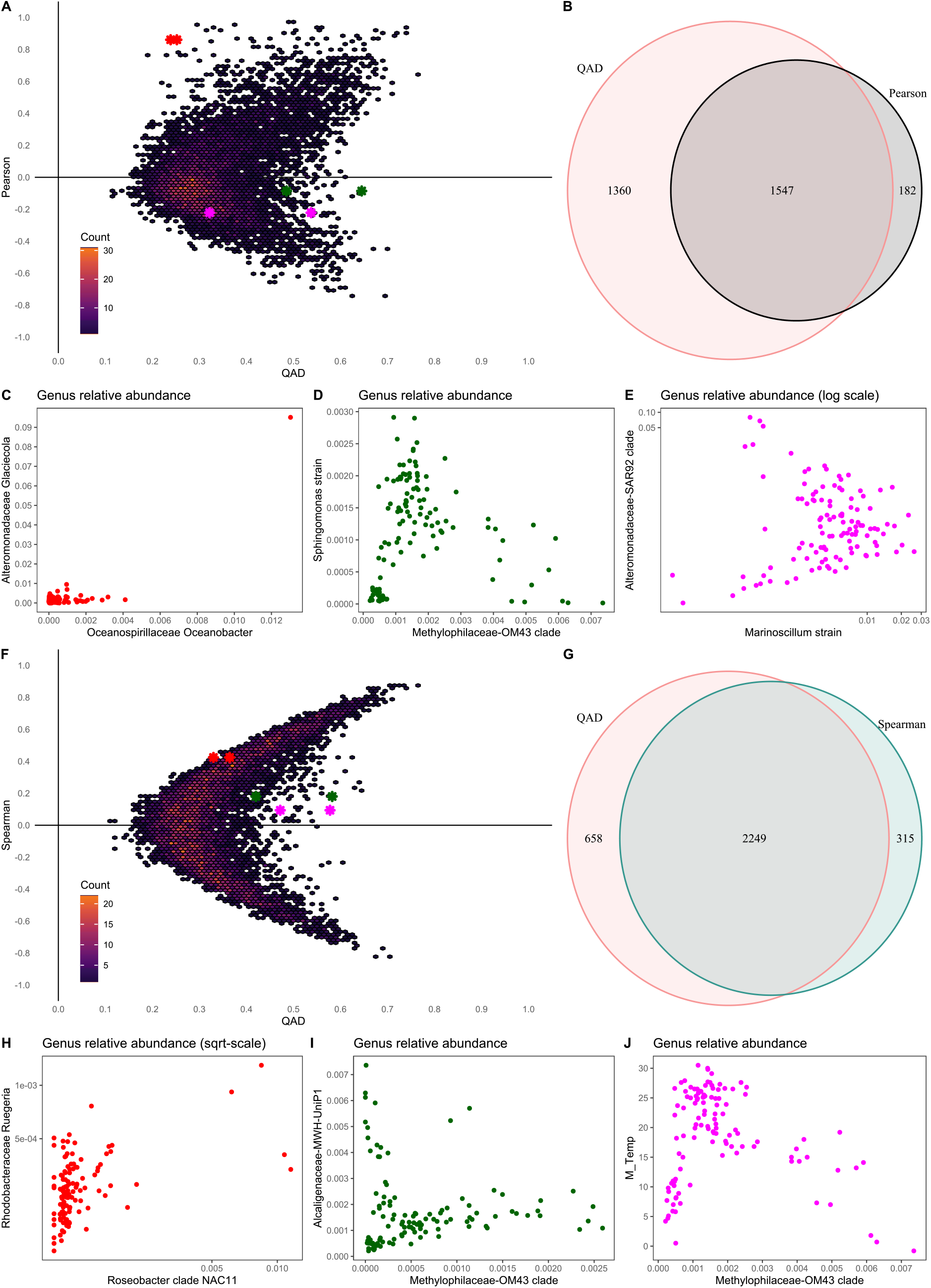
Application of Pearson’s *r*, Spearman’s *ρ* and *qad* to a subset of the Tara Oceans dataset. **(A, F)** Hex bin plot of qad vs. Pearson’s *r* (Spearman’s *ρ*) for all pairwise relationships; colour code corresponding to count numbers per hexagonal bin. **(B, G)** Venn diagrams depicting the number of significant relationships across all pairwise associations. **(C**,**H)** Scatterplots of selected pairwise relationships significant w.r.t. Pearson’s *r* (Spearman’s *ρ*) but not w.r.t. *qad*. **(D**,**E**,**I**,**J)** Scatterplots of selected pairwise relationships (highly asymmetric) which are not significant w.r.t. Pearson’s *r* (Spearman’s *ρ*) but highly significant w.r.t. the measure *qad*. The colour of the points corresponds to the coloured stars in the hex bin plot depicting the *q*-value as well as the Pearson (Spearman) correlation.

## CONCLUSION

Our theoretical and real-world examples demonstrate that the measure *qad* (as well as the measures mentioned in Table 1) is able to quantify and indicate the extent of dependence in general situations, whereas classical measures only capture linear and monotonic associations. In most real-world situations no, or almost no, prior knowledge about the interdependence of variables is available. Thus, aiming at an objective estimate of the strength of (directed) dependence it is therefore unavoidable to work with measures not relying on distributional assumptions. Considering non-monotonic and non-functional relationships naturally expands our ability to detect more complex, and potentially asymmetric relationships between organisms and their environment. We demonstrated that neither of the methods discussed here outperforms all other methods in full generality, every statistical tool exhibits limitations in specific settings. If it is known in advance that the data originates from a linear or a monotonic setting, we recommend classical measures of association such as Pearson’s *r* and Spearman’s *ρ*. In most situations (e.g., in network inference, data mining), however, wrongly imposing linearity/monotonicity without prior knowledge may lead to wrong conclusions. In general cases we therefore strongly recommend the use of more recent methods for quantifying pairwise dependencies. We showed that *qad* is powerful in detecting dependence and provides reliable and easily interpretable results.

Another important property of bivariate associations is asymmetry and direction in dependence in the sense that the information gain/predictability of quantity Y given knowledge of quantity X is not the same as *vice versa*. Considering direction and asymmetry in dependence facilitates the detection and extraction of patterns from ecological datasets and the testing of refined hypotheses. For instance, correlation analysis testing for relationships between the abundance of pairs of taxa is usually performed as basis for network inference, which, in turn, facilitates the interpretation of microbiome structure. Ecological relationships between organisms may be reciprocal in the sense that taxa mutually affect each other, either positively (mutualism) or negatively (competition). They may, however, also be directed in such a way that a given taxon is facilitating or inhibiting the growth of another taxon without being affected itself by the other taxon (e.g., commensalism, amensalism). As shown before, conventional correlation analysis neither detects directed relationships nor discriminates between directed and mutual relationships and is therefore of limited value for the interpretation of community dynamics. We are aware of only two non-parametric methods that are able to quantify directed dependence, namely *qad* (Junker, Griessenberger & Trutschnig 2021) and *xicor (Chatterjee 2020)*. We have shown that *qad* has a higher overall power in detecting deviation from independence, especially in very noisy data sets *qad* seems to perform better than *xicor*. Furthermore, the R-package *qad* provides use-friendly outputs and a number of additional features that facilitate the interpretation of the results as well as functions to use *qad* as a prediction tool.

We conclude that the interpretation of ecological data may be strongly biased by the choice of statistical approaches quantifying dependence between two random variables. The acknowledgement and adequate handling of asymmetry, a universal property of bivariate associations, is an important step towards additional information gain and the avoidance of model bias for small, medium and large datasets, and will advance and allow for a deeper understanding of ecological systems.

## Acknowledgments

This study was funded by the Austrian Science Fund (FWF, Y 1102 B29) granted to RRJ. Moreover, the first and the second author gratefully acknowledge the support of the WISS 2025 project ‘IDA-lab Salzburg’ (20204-WISS/225/197-2019 and 20102-F1901166-KZP).

## Author contributions

FG, RRJ and WT designed the study; FG analysed the data; FG, RRJ and WT wrote the manuscript.

## Competing interests

The authors declare no competing financial interests.

## Data availability statement

All data used in the study can be found at other sources (mentioned at the corresponding paragraphs) or in the Supplementary information.

## Code availability

The *qad* package is available for the R programming language and can be downloaded at https://cran.r-project.org/web/packages/qad/index.html. This paper describes the latest CRAN-version of *qad* (v.1.0.1). To install the package, run *install*.*packages(‘qad’)*. The development version of *qad* is available on GitHub (https://github.com/griefl/qad) and can be installed by running *devtools::install_github(“griefl/qad”, dependencies = TRUE, build_vignettes = TRUE)*.

## Supplementary Information

Supplementary information 1: R-Code (Supplementary information 1.R) including data generation from the studied dependence structures for the empirical experiments on extreme settings and power analysis.

Supplementary information 2: Additional R-Code and shiny app (dep_measures.R and app.R) for studying the empirical behaviour of the eight dependence measures on various dependence structures, noise levels and sample sizes.

Supplementary information 3: Additional results/plots of the comparison with other dependence measures (dependence scores in extreme settings, empirical power analysis and runtime analysis).

## Notes

### Competing Interest Statement

The authors have declared no competing interest.

