## SupplementaryInformation for "*qad:* An R-package to detect asymmetric and directed dependence in bivariate samples": Supplementary Information3.pdf

### Supplementary information 3

#### Performance and comparison of qad with other dependence measures

In order to compare the performance of qad with the seven dependence measures listed in Table 1 (in the main paper) in various dependence settings we generated data points from nine different dependence structures with increasing noise (see Supplementary Information 1). Noise was generated by drawing random numbers from a uniformly distributed random variable on the interval $[-a, a]$, whereby $a$ was increased from 0 to 1 by steps of 1/100. We repeated the experiment R=1.000 times. Dependence measures such as qad that capture the dependence in general situations should assign similar scores of dependence to equally noisy data independently of the concrete functional relationship (Reshef et al. 2011). Accordingly, the measures of dependence capable of detecting functional dependence, i.e., MIC, qad, RCD, rdc and xicor showed a similar decrease in precision with increasing noise irrespective of the functional relationship between X and Y (see Fig. 3 A,B,D,E,F,I in the main paper). On the other hand, Pearson's r and Spearman's ρ failed to share this property; in the linear setting, for instance, the values of Pearson's r and Spearman's ρ decreased with increasing noise, whereas in the context of a parabola/torus or cross both measures returned an average value of 0 irrespectively of the noise (Fig. 3 A,B,G and H in the main paper). Furthermore, in the linear setting the quantities dCor and rdc returned dependence values close to Pearson's r and Spearman's ρ followed by qad. The other measures (MIC, xicor and RCD) deviated from Pearson's r and Spearman's ρ to a larger extent (Fig. 3A in the main paper).

Considering the opposite extreme (i.e., independence) the estimated dependence measures showed a slightly different behavior. The methods xicor, Pearson's r and Spearman's ρ spread around 0, whereas the other dependence measures were strictly positive (Fig. 3C in the main paper). The reason for this behavior is that Pearson's r, Spearman's ρ and xicor can attain negative values too. In the case of Pearson's r and Spearman's ρ, negative values are natural and can be interpreted accordingly. In the case of xicor, which is an estimator of a theoretical dependence measure with values between 0 and 1, negative values are very difficult, if not impossible to interpret appropriately for applicants. Notice that for dependence measures which are strictly positive (e.g., qad), deviation from 0 in the case of independence can be ignored for very large sample sizes (n >> 1000), in small sample settings, however, the deviation is essential for interpreting the obtained dependence scores. As example, assume we have a bivariate sample of size n=100 drawn from independent random variables X and Y, and a calculated dependence value of qad(X,Y)=0.2. Then this qad value points towards independence, which is also reflected by a p-value > 0.05. Only focusing on the obtained value is clearly insufficient for deciding if, or if not, the sample is likely to come from independent random variables – variability of the obtained values has to be taken into consideration. Overcoming this problem, qad calculates a p-value coinciding with the probability that a q-value - calculated from a sample of independent random variables - is greater than or equal to the actual q-value of the sample, this allows to interpret the obtained values and puts them into perspective.

Furthermore, the measures qad and xicor, which detect asymmetry in dependence by construction, recognized asymmetry in settings in which (on average) more information on Y could be obtained by knowing the value of X than vice versa, i.e., qad(X,Y)>qad(Y,X) as well as xicor(X,Y)>xicor(Y,X), see, for instance, Fig. 3 B,D,E,F in the main paper.

**Statistical power analysis**

In order to compare the statistical power for detecting deviations from independence we followed (Simon & Tibshirani 2014) and generated bivariate samples drawn from functional and non-functional relationships (see Supplementary information 1) with respect to eight different levels of vertically added noise (normally distributed with mean 0 and standard deviation $\sigma \in (0, 0.1, 0.25, 0.5, 0.75, 1, 1.25, 1.5)$) and increasing sample size (from $n = 10$ to $n = 500$ with increments of $10$). To determine a threshold for testing independence we generated independent data with the same marginal distributions of the previously defined dependence structures (see Supplementary information 1) and estimated the empirical distribution for each of the methods with $R = 1000$ runs. As cut-off value we determined the 0.95-quantile, respectively. Hence, the statistical power was computed as fraction of the 1000 computed dependence values which were greater than the respective cut-off value (empirical p.value).

Statistical power (for detecting that the random variables are not independent) varied strongly across the measures for different dependence structures (Fig. 1). We found that no dependence measure globally dominated all other measures, i.e., different dependence structures were detected with different power by different methods. For instance, in the linear setting (Fig. 1A) or in the setting with a monotonic trend (on average), e.g. “Two lines” (Fig. 1C), the measures of *Pearson*, *Spearman* and *dCor* had very high power compared to *qad*, *rdc* and *MIC$_e$*, which, in turn, had higher power than *xicor* and *RCD.* In non-linear and non-monotonic settings (e.g., sinusoid, parabola – Fig. 1B, D) the methods capable of detecting non-monotonic relationships outperformed *Pearson* and *Spearman correlation* (which, in this case, might completely fail to detect any deviation from independence). The results were consistent for different choices of noise levels and other dependence structures (see the Figures in the following). Considering the normalized average area under the power curves as an indicator for good overall performance to detect deviation from independence, we found that the dependence measures *qad* and *MIC$_e$* were the best performing methods with respect to the minimum average area under the power curve, followed by *xicor*, *RCD* and *rdc* (Fig. 1E).

Focusing on the median overall power, the measures *dCor, MICe,* rdc and *qad* showed very high and similar values and can, therefore, be recommended over *Pearson's r, Spearman's* ρ*, RDC* and *xicor* in general settings. If information on linear or monotonic associations is desired, dependence measures should be chosen accordingly. Furthermore, commonly used methods such as *Pearson's* $r$ or *Spearman's* ρ showed a lot of variability in statistical power, which makes them less applicable in general settings, where the underlying relationship is completely unknown. Altogether, our results were mostly consistent with previous findings of power analyses on a subset of the selected dependence measures (Lopez-Paz, Hennig & Schölkopf 2013; Reshef *et al.* 2018). As a consequence, we conclude that the commonly used measures *Pearson's* $r$ and *Spearman's* $\rho$ have very low power in a variety of dependence settings and are therefore not the optimal choice for quantifying dependence without prior knowledge on the data's underlying dependence structure (e.g., linearity assumption in the case of *Pearson's r, or monotonicity in the case of Spearman's* ρ). The measures *qad* and $MIC_e$ seem to be applicable in general settings since they exhibited reasonable power (in detecting dependence) in each of the tested settings. Without prior knowledge it is recommendable to work with one of these two approaches.

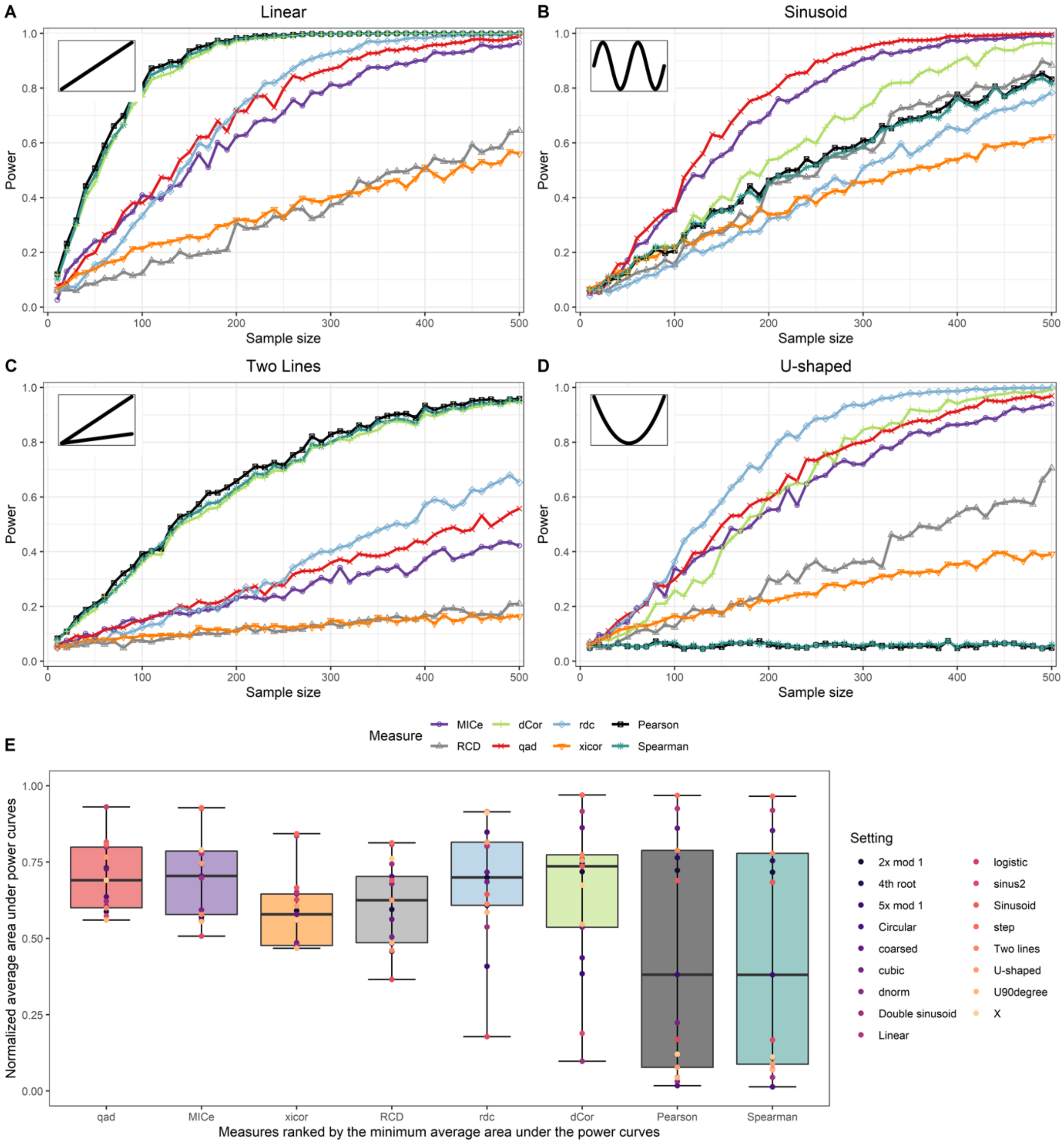


Figure 1. **(A-D)** Subset of four (slightly noisy) relationships considered in statistical power analysis. Empirical power is illustrated for a relationship with vertically added noise following a normal distribution with mean 0 and standard deviation 1. The underlying noise-free relationship is depicted in the left top corner. In each setting, the sample size increases from left (n =10) to right (n = 500) with increments of 10. In order to estimate the power, 1000 runs were executed for each sample size **(E)** Normalized average area under the power curves for the measures *qad, MICe, rdc, xicor, RCD, dCor, Pearson* and *Spearman* correlation for all 17 dependence structures. The boxplots are ordered by the minimum average area under the power curve.

**Runtime Analysis**

In the era of Big Data runtime of the implemented methods is substantial. Therefore, we recorded the average runtimes for each method. More precisely, we generated bivariate samples drawn from independent uniformly distributed random variables on $[0,1]$ and computed the runtime for one evaluation (without p-values) for increasing sample sizes $n \in (10,100,1.000,10.000,100.000)$. We repeated the experiment 100 times and averaged the obtained values. Since for distance correlation and RCD the memory allocation failed for a sample of size $n = 100.000$ or the evaluation time exceeded 5 minutes, we did not get values of these settings.

Overall, the runtime performance of *Pearson's r, qad, rdc*, *Spearman's* $\rho$ and *xicor* was quite similar, these methods outperformed the others when it comes to runtime for very large sample sizes (Table 1). The measures *dCor,* $MIC_e$ and *RCD* were more costly to calculate, the running time was therefore higher. The performance of *RCD* and *dCor* was acceptable for sample sizes below $n = 10{,}000$, but extended the predefined threshold for a sample of size $n = 100{,}000$.

| n | dCor | $MIC_e$ | Pearson | qad | RCD | rdc | Spearman | xicor |
|---|---|---|---|---|---|---|---|---|
| 10 | <0.01 | <0.01 | <0.01 | <0.01 | <0.01 | <0.01 | <0.01 | <0.01 |
| 100 | <0.01 | <0.01 | <0.01 | <0.01 | 0.01 | <0.01 | <0.01 | <0.01 |
| 1.000 | 0.15 | 0.17 | <0.01 | <0.01 | 0.22 | <0.01 | <0.01 | <0.01 |
| 10.000 | 14.71 | 3.79 | <0.01 | <0.01 | 15.55 | 0.03 | <0.01 | <0.01 |
| 100.000 | NA | 79.04 | <0.01 | 0.06 | NA | 0.31 | 0.03 | 0.04 |

NA: The evaluation time exceeded 5 minutes, or the memory allocation failed, respectively.

Table 1: Average run times (in seconds) for one calculation given a bivariate data sample of size $n \in (10,100,1000,10000,100000)$.

### Additional Figures of the power analysis

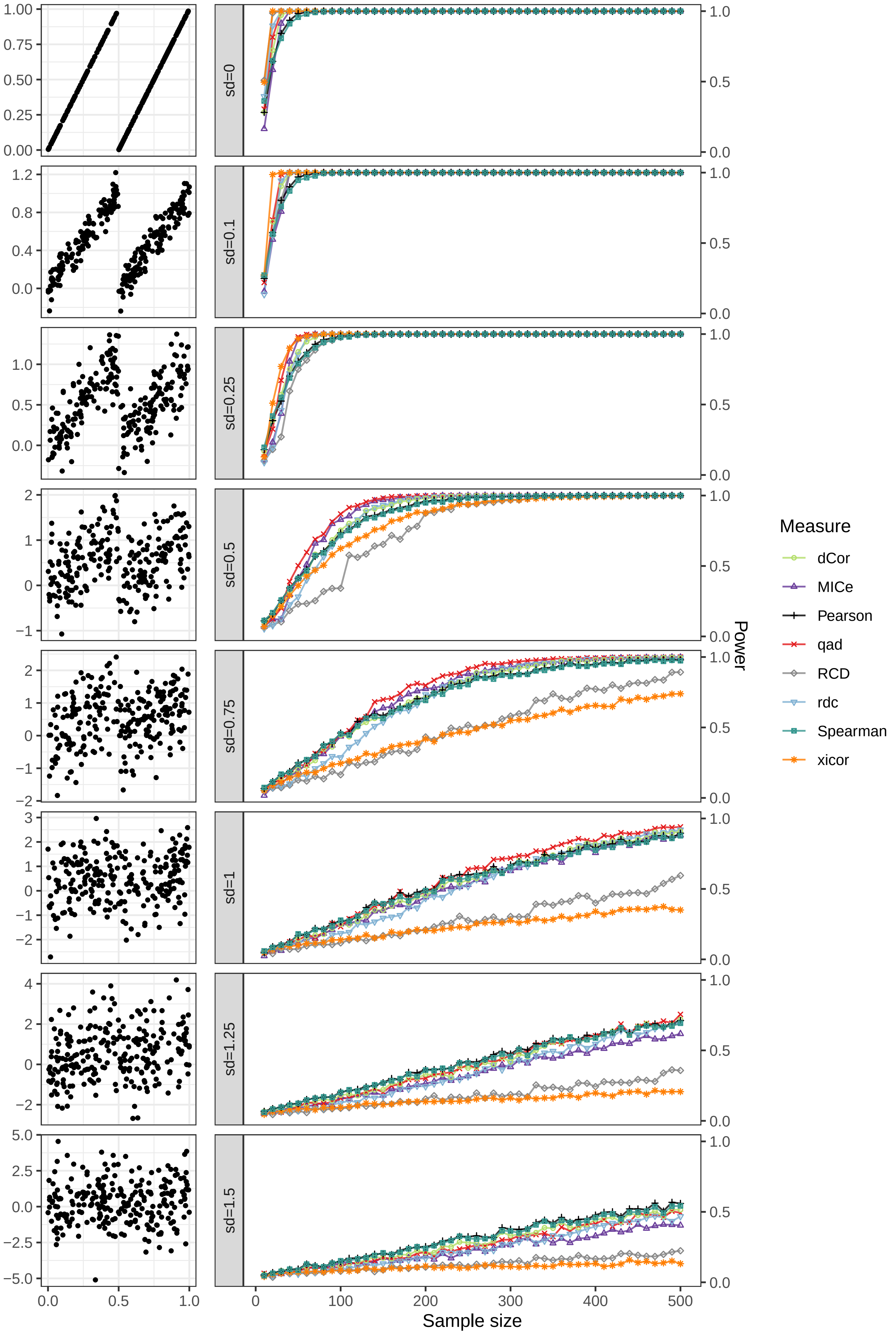

sd=0
sd=0.1
sd=0.25
sd=0.5
sd=0.75
sd=1
sd=1.25
sd=1.5
Sample size
Power
Measure
dCor
MICe
Pearson
qad
RCD
rdc
Spearman
xicor

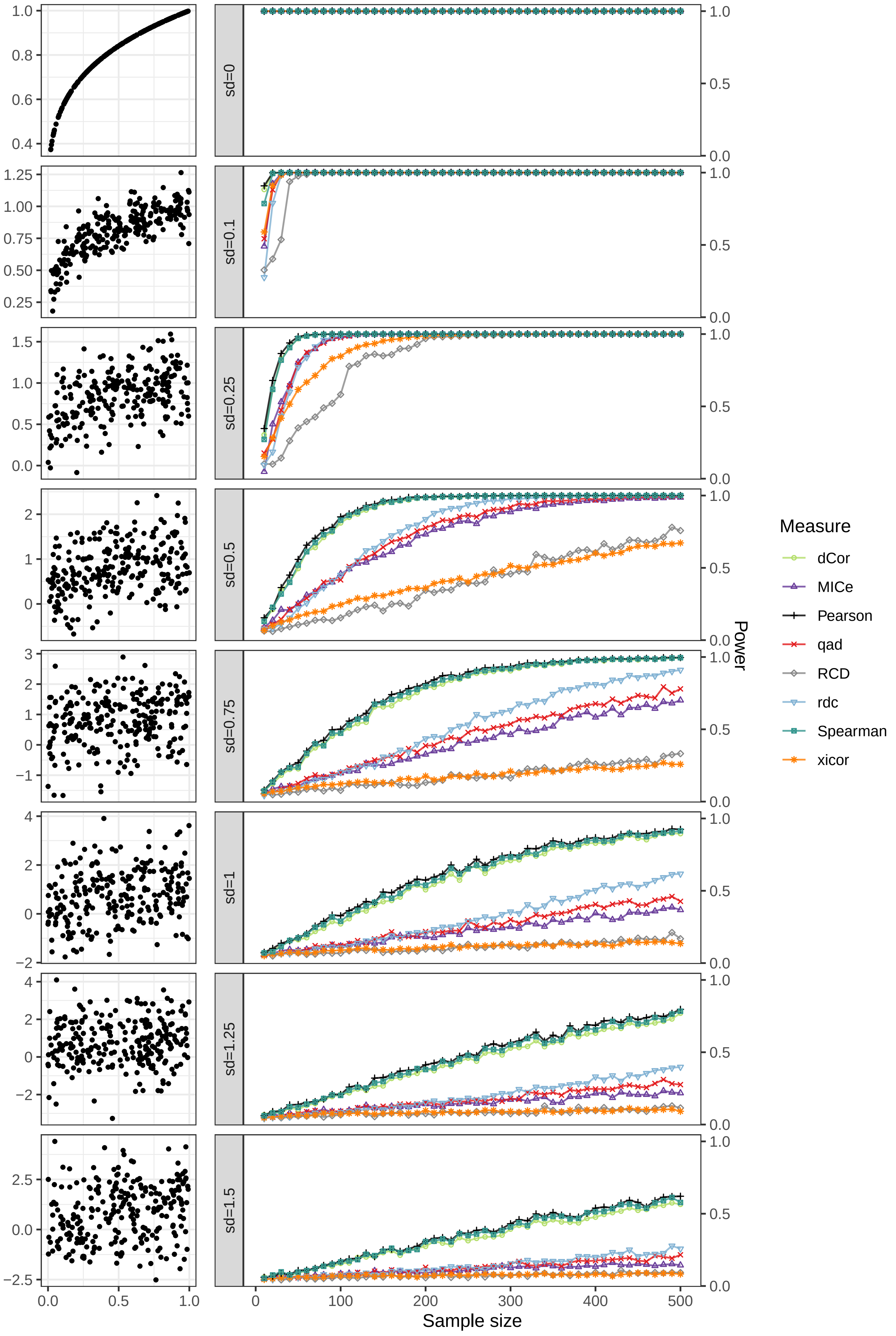

sd=0
sd=0.1
sd=0.25
sd=0.5
sd=0.75
sd=1
sd=1.25
sd=1.5
Power
Sample size
Measure
dCor
MICe
Pearson
qad
RCD
rdc
Spearman
xicor

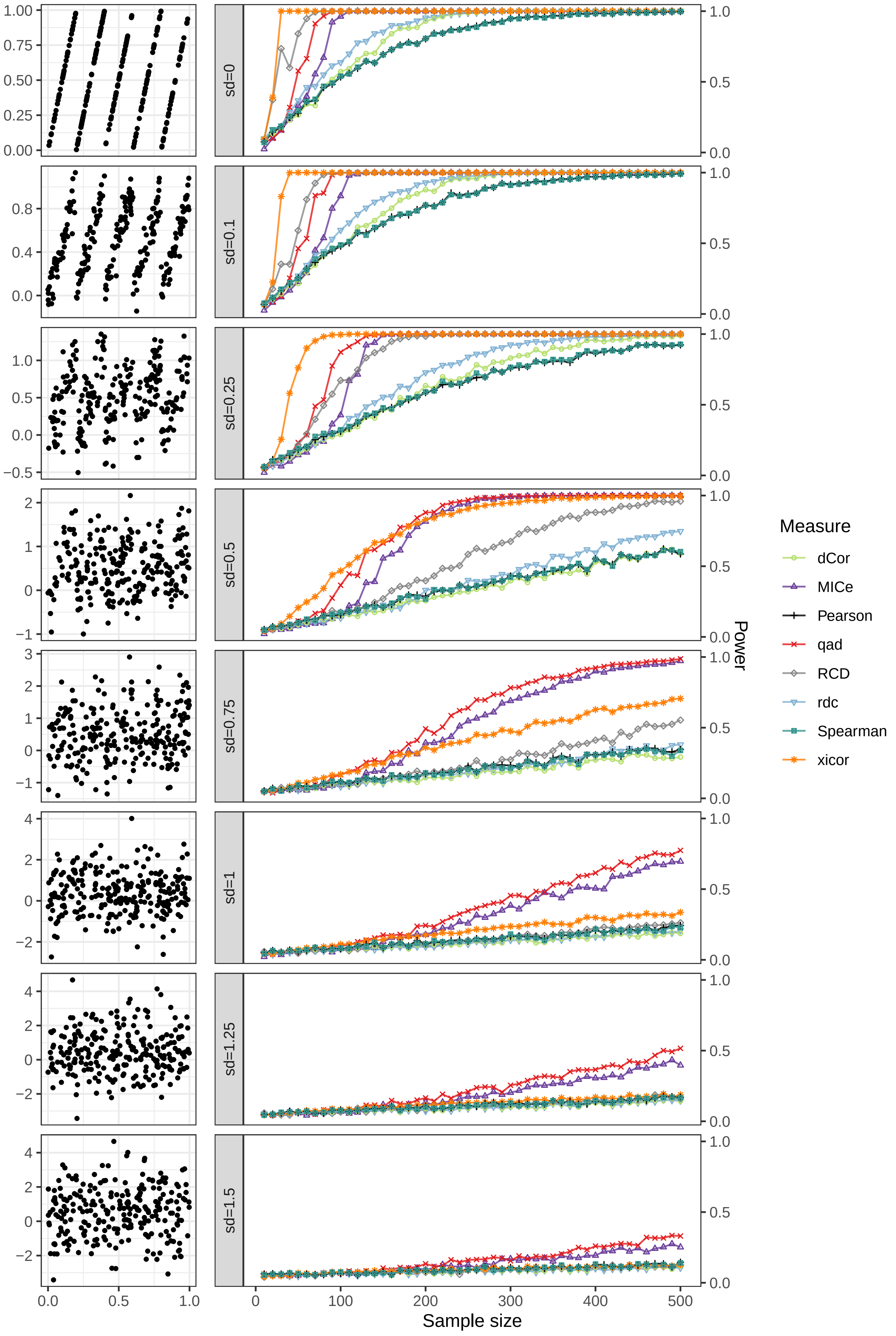

sd=0
sd=0.1
sd=0.25
sd=0.5
sd=0.75
sd=1
sd=1.25
sd=1.5
Sample size
Power
Measure
dCor
MICe
Pearson
qad
RCD
rdc
Spearman
xicor

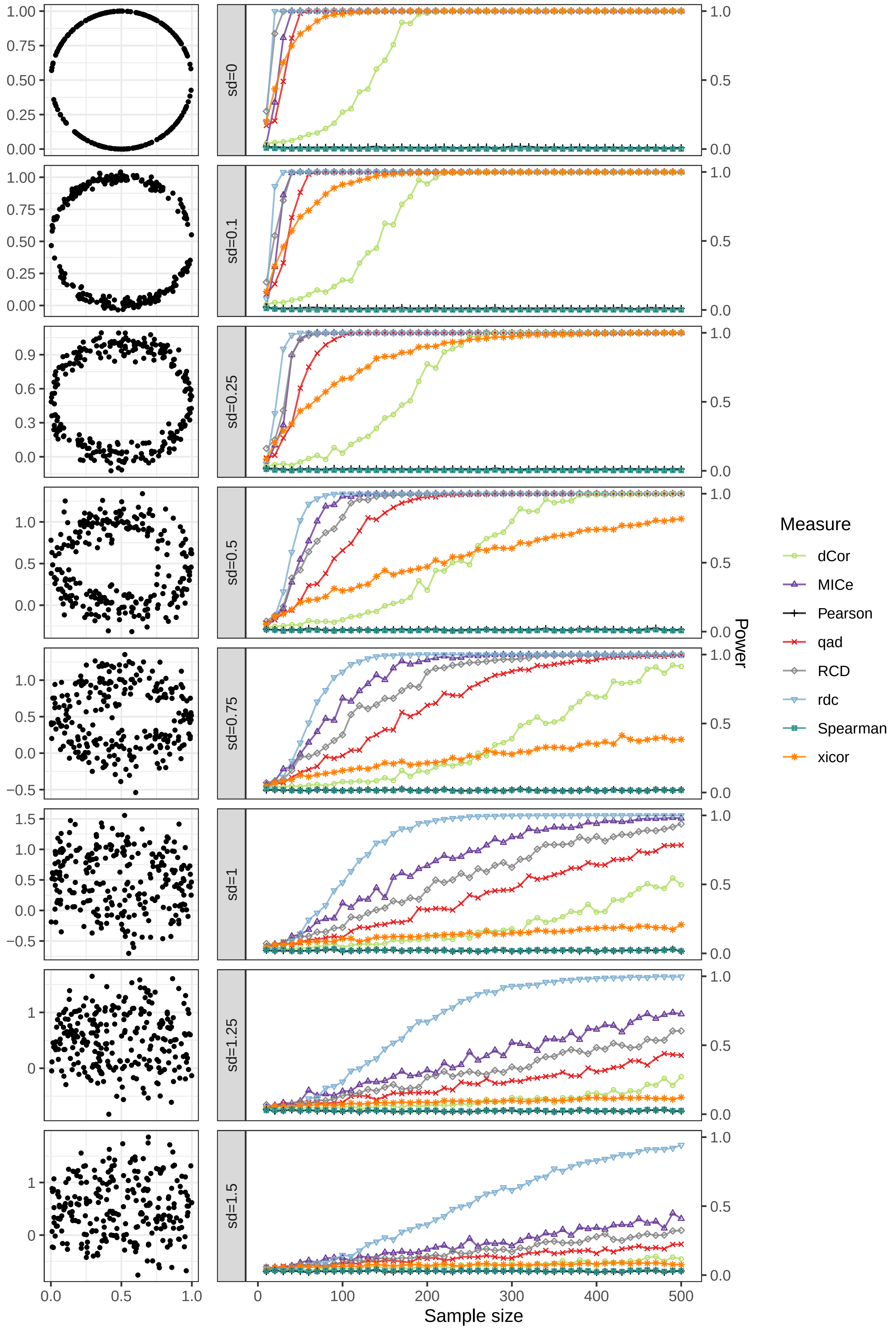

sd=0
sd=0.1
sd=0.25
sd=0.5
sd=0.75
sd=1
sd=1.25
sd=1.5
Sample size
Power
Measure
dCor
MICe
Pearson
qad
RCD
rdc
Spearman
xicor

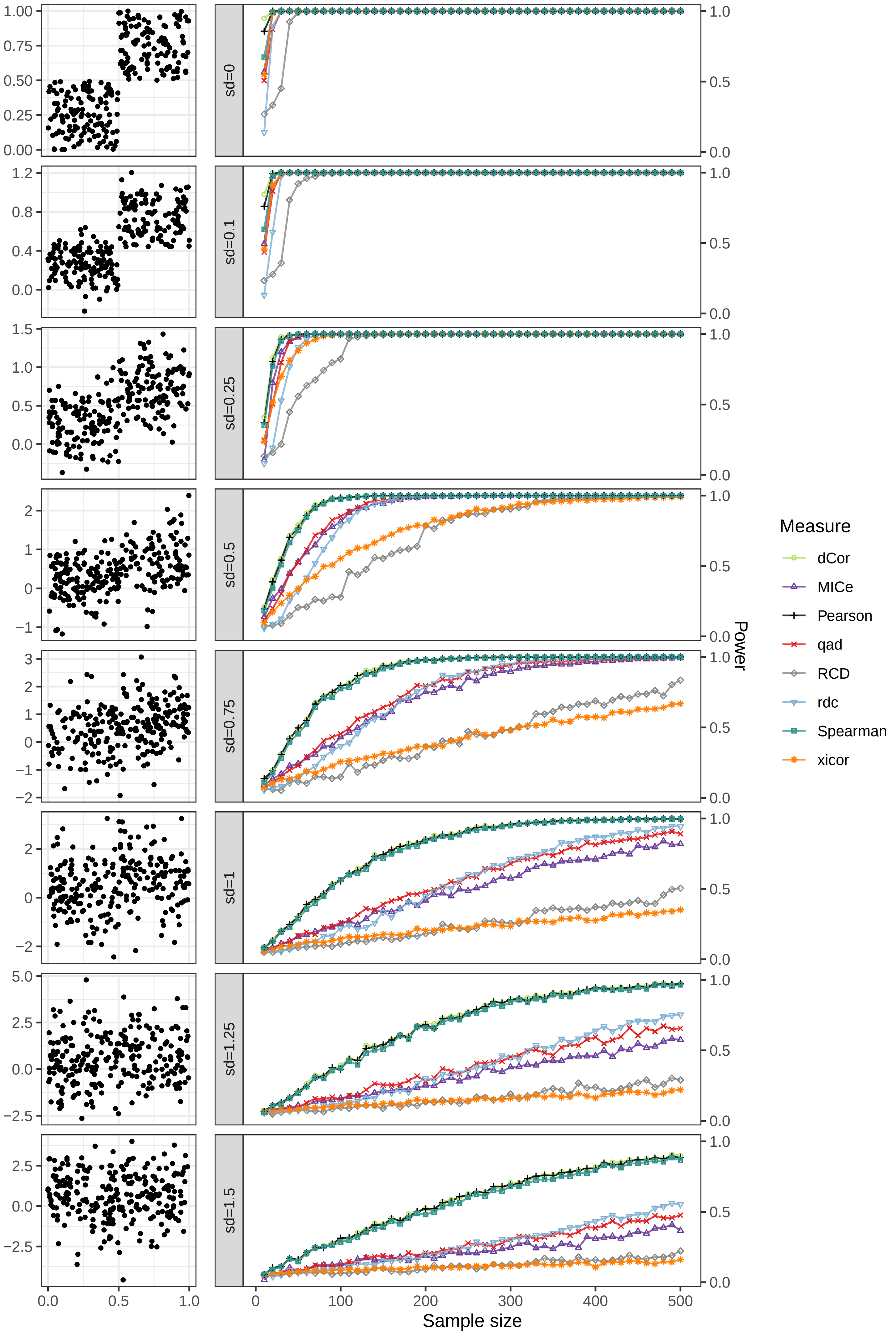
sd=0
sd=0.1
sd=0.25
sd=0.5
sd=0.75
sd=1
sd=1.25
sd=1.5
Measure
dCor
MICe
Pearson
qad
RCD
rdc
Spearman
xicor
Power
Sample size

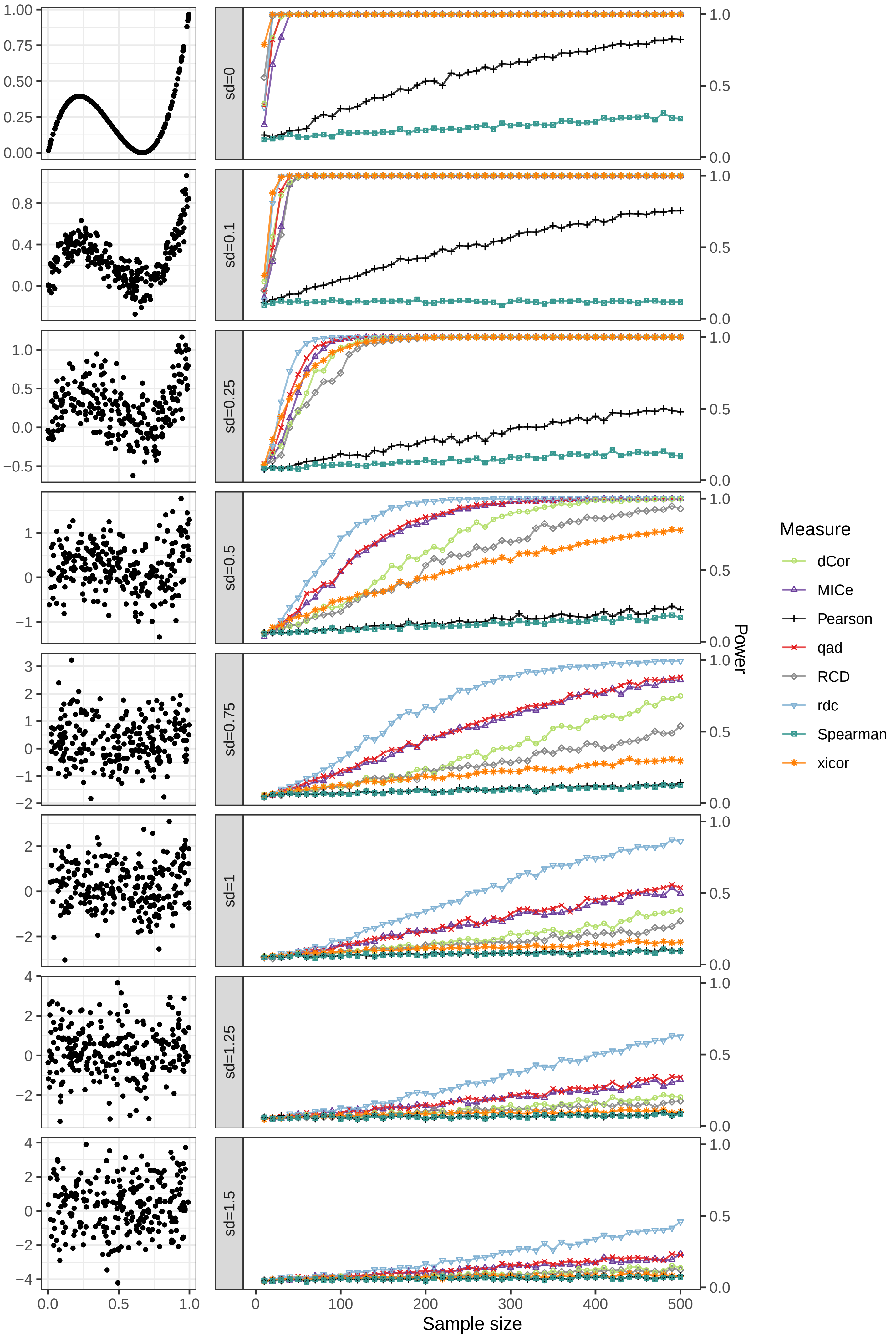
sd=0
sd=0.1
sd=0.25
sd=0.5
sd=0.75
sd=1
sd=1.25
sd=1.5
Sample size
Power
Measure
dCor
MICe
Pearson
qad
RCD
rdc
Spearman
xicor

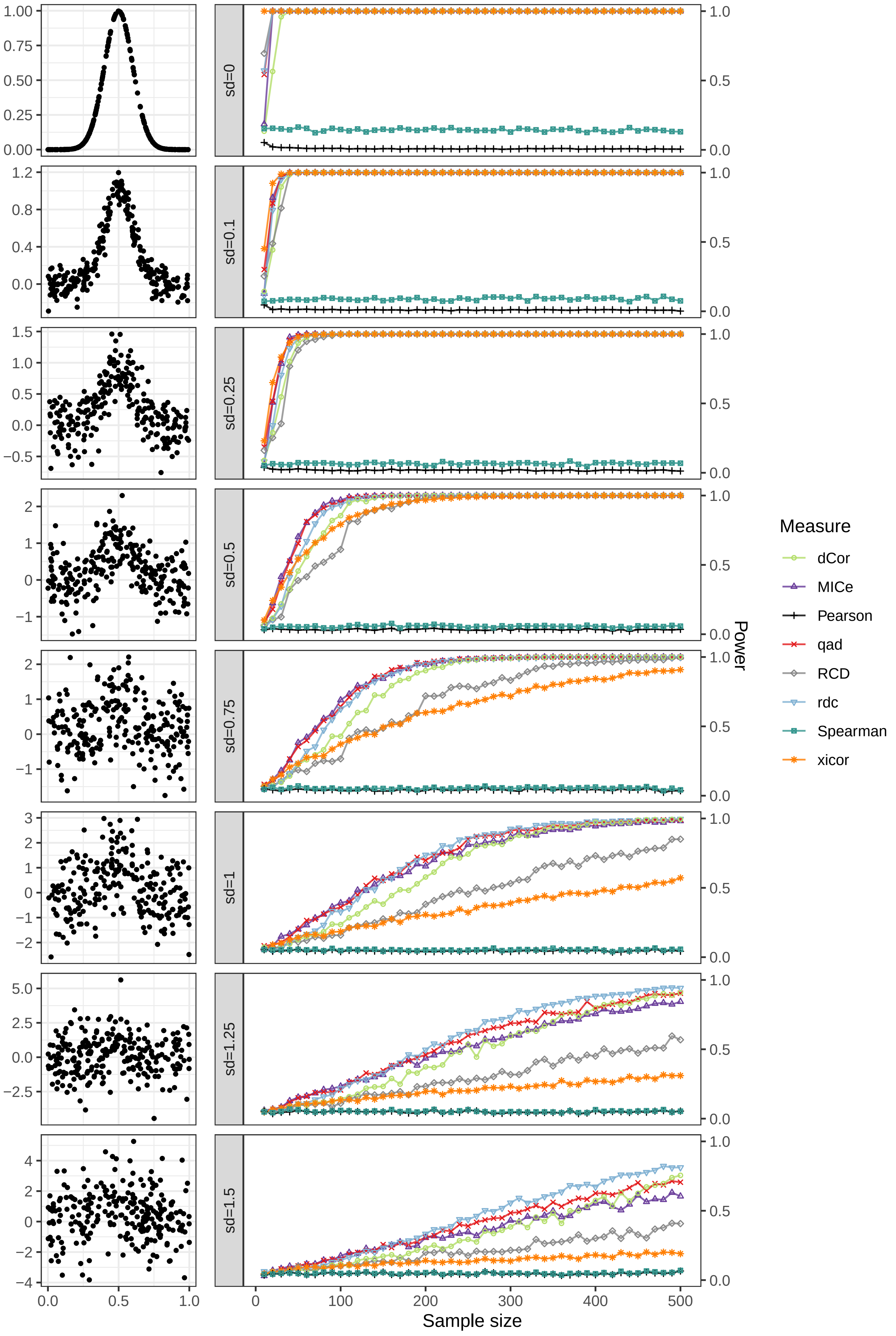
sd=0
sd=0.1
sd=0.25
sd=0.5
sd=0.75
sd=1
sd=1.25
sd=1.5
Sample size
Power
Measure
dCor
MICe
Pearson
qad
RCD
rdc
Spearman
xicor

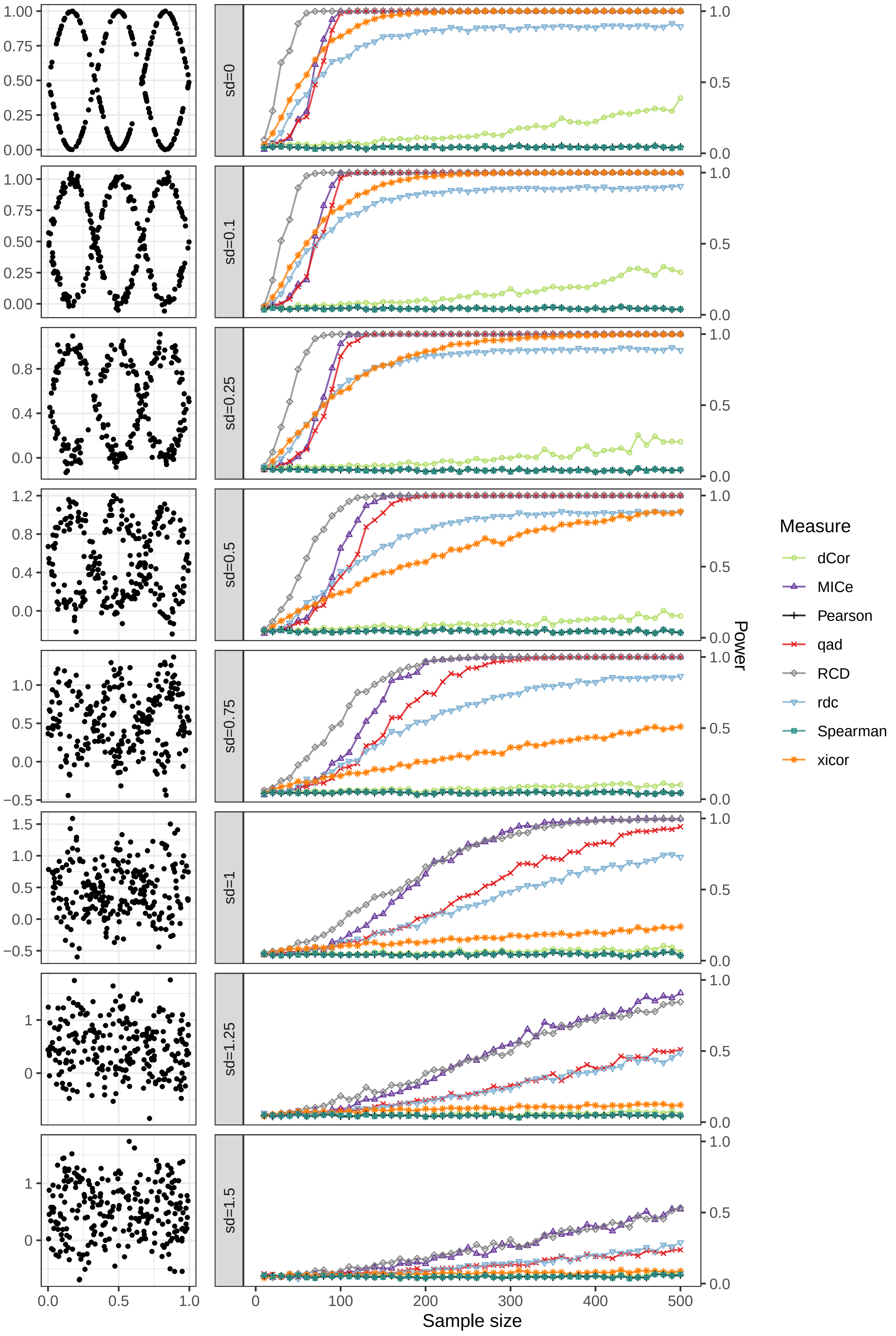

sd=0
sd=0.1
sd=0.25
sd=0.5
sd=0.75
sd=1
sd=1.25
sd=1.5
Measure
dCor
MICe
Pearson
qad
RCD
rdc
Spearman
xicor
Power
Sample size

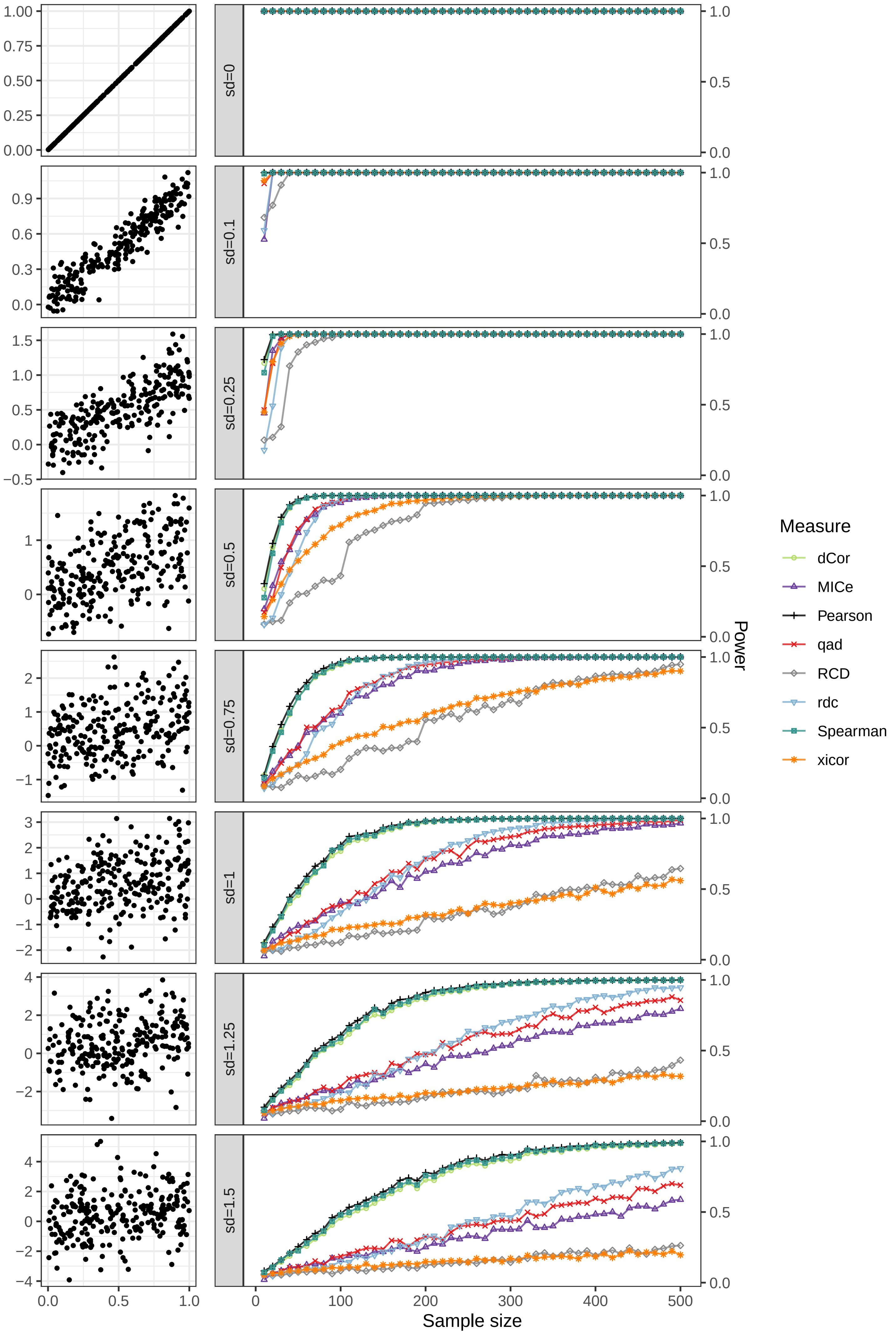

sd=0
sd=0.1
sd=0.25
sd=0.5
sd=0.75
sd=1
sd=1.25
sd=1.5
Sample size
Power
Measure
dCor
MICe
Pearson
qad
RCD
rdc
Spearman
xicor

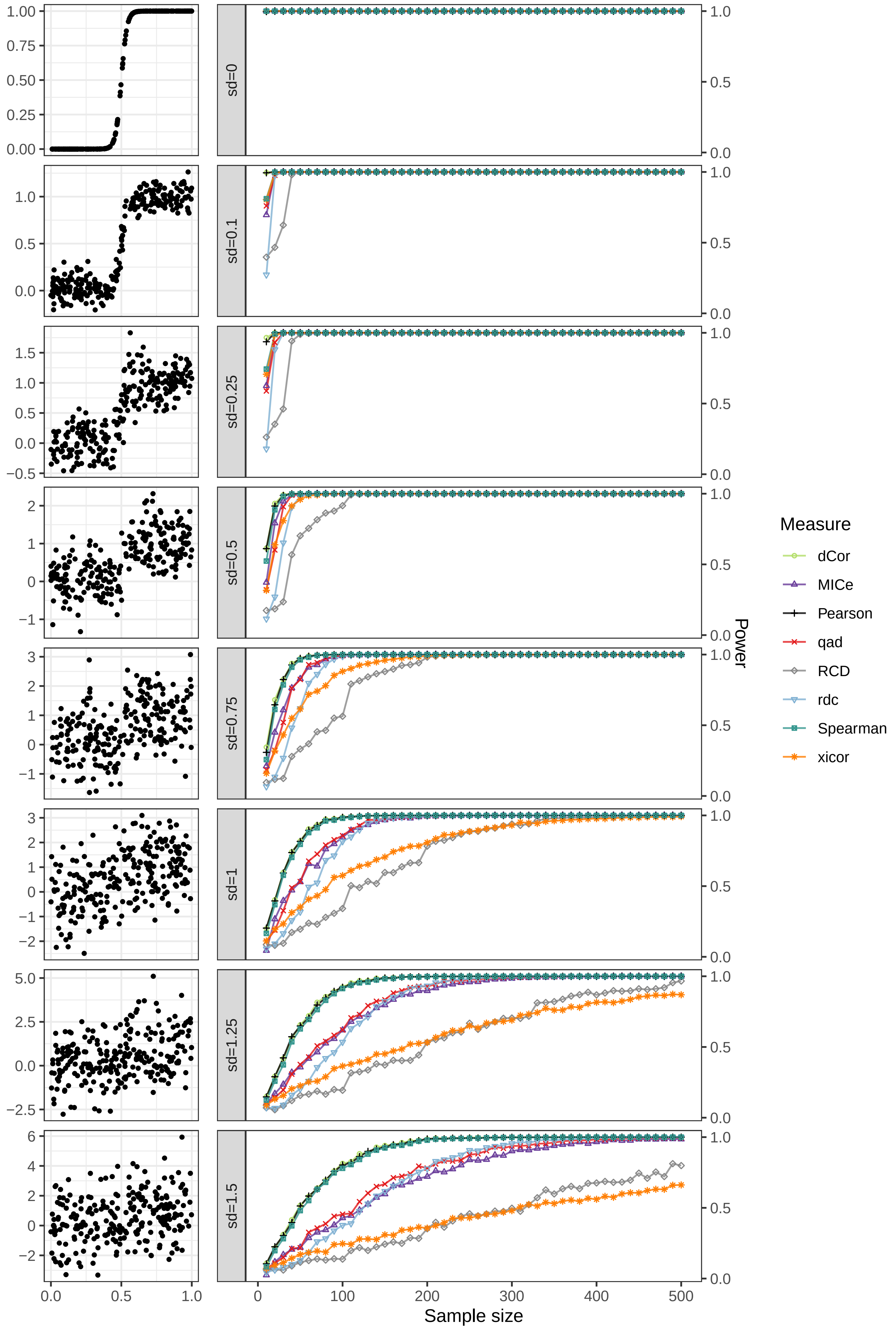

sd=0
sd=0.1
sd=0.25
sd=0.5
sd=0.75
sd=1
sd=1.25
sd=1.5
Sample size
Power
Measure
dCor
MICe
Pearson
qad
RCD
rdc
Spearman
xicor

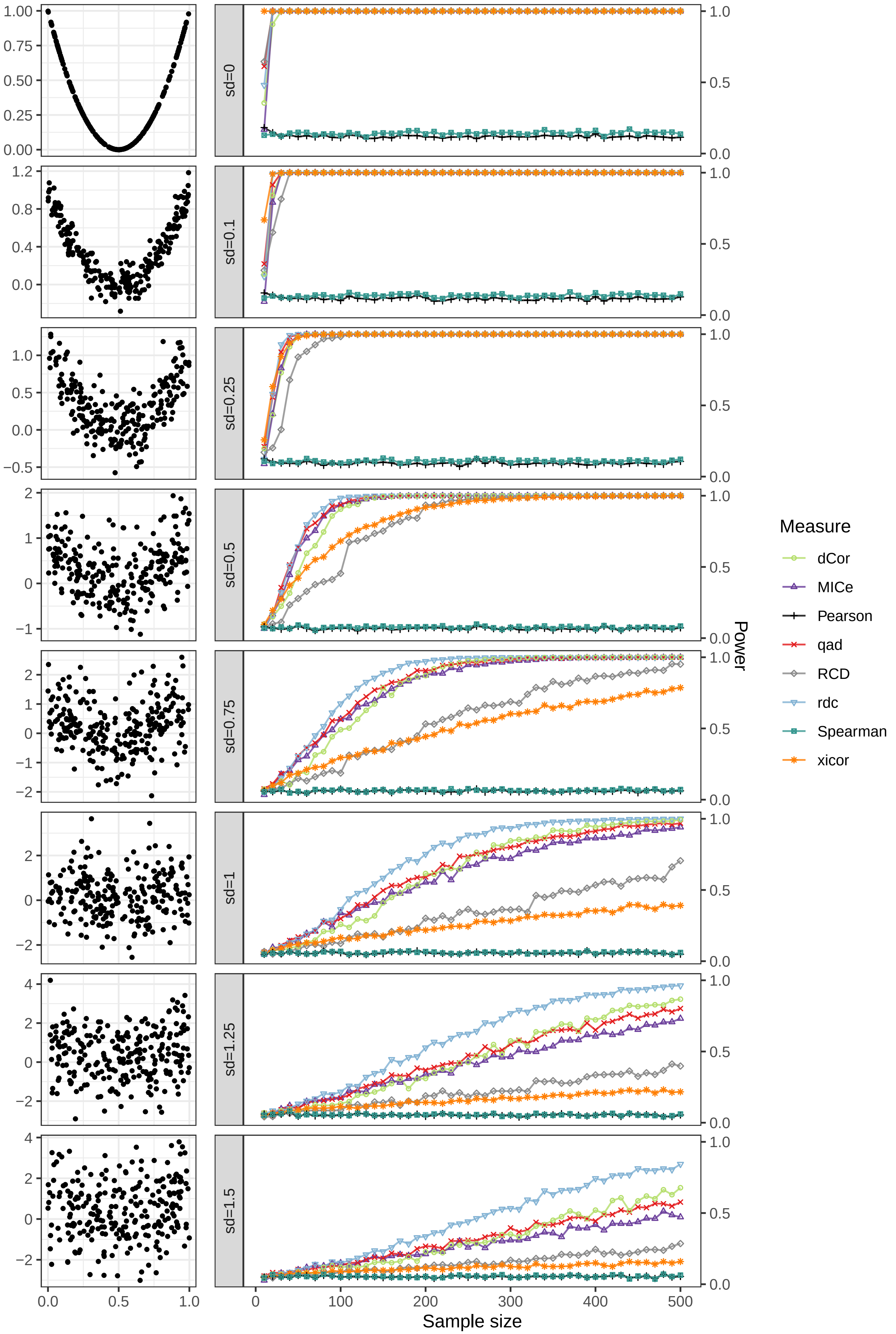
sd=0
sd=0.1
sd=0.25
sd=0.5
sd=0.75
sd=1
sd=1.25
sd=1.5
Sample size
Power
Measure
dCor
MICe
Pearson
qad
RCD
rdc
Spearman
xicor

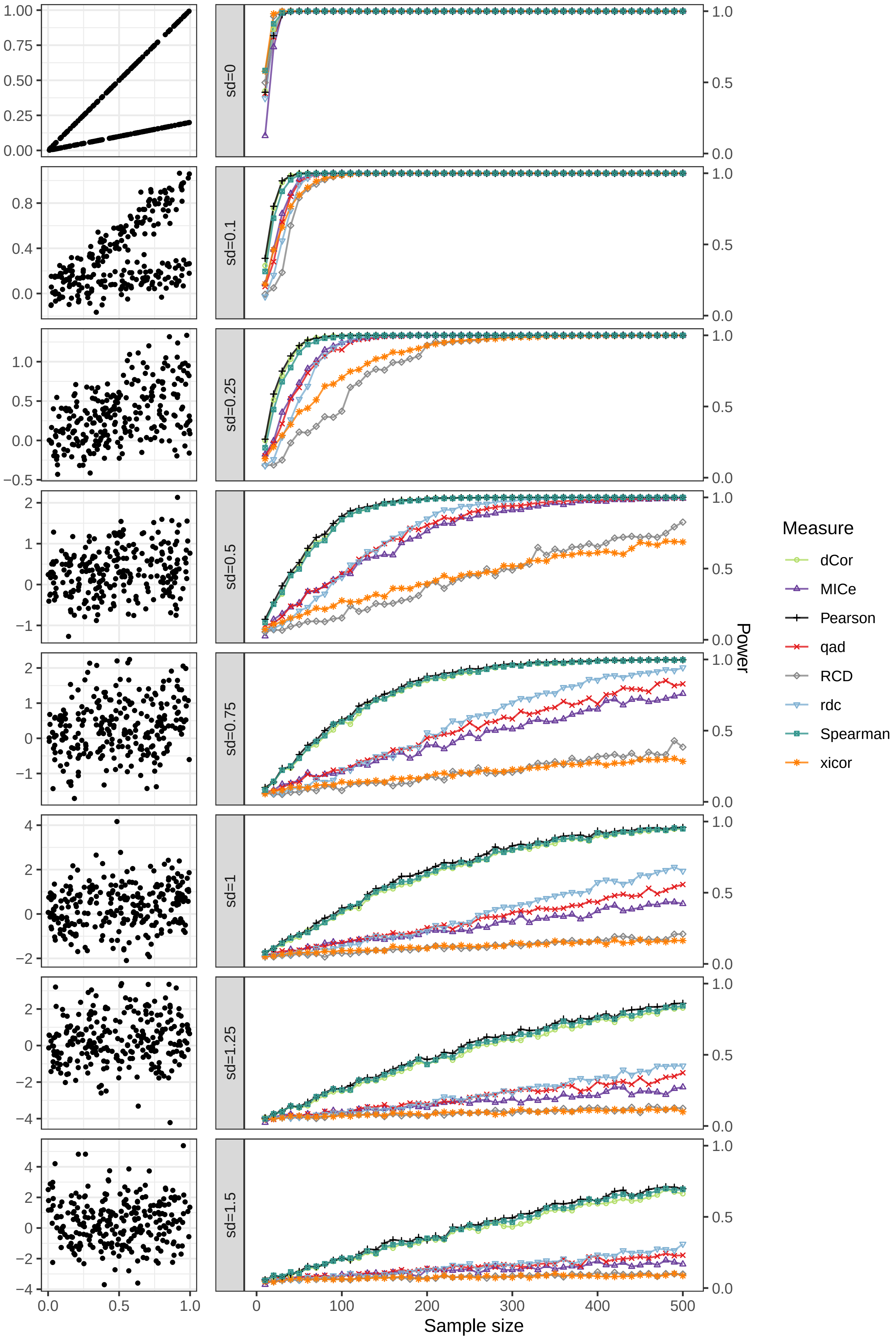

sd=0
sd=0.1
sd=0.25
sd=0.5
sd=0.75
sd=1
sd=1.25
sd=1.5
Sample size
Power
Measure
dCor
MICe
Pearson
qad
RCD
rdc
Spearman
xicor

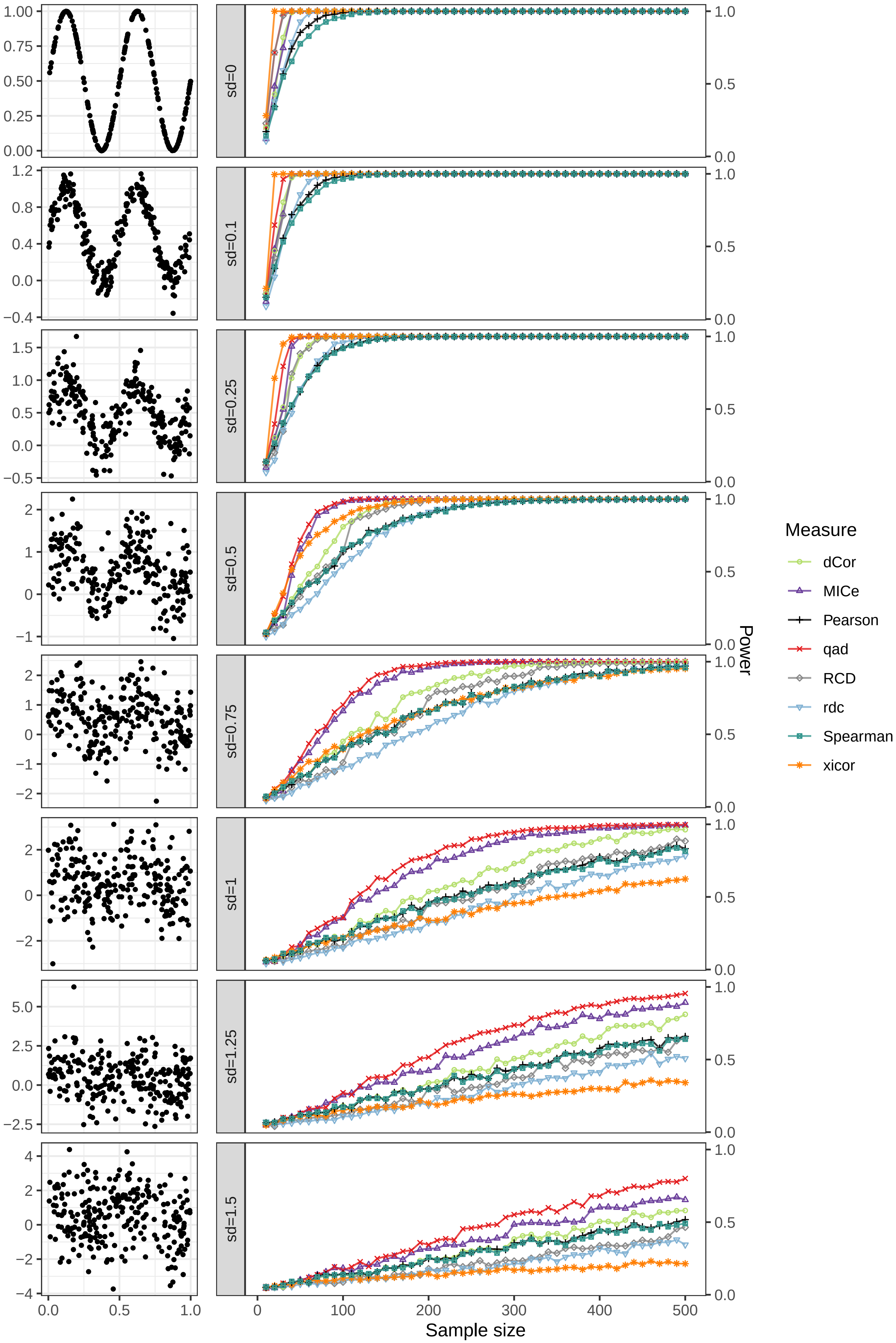

sd=0
sd=0.1
sd=0.25
sd=0.5
sd=0.75
sd=1
sd=1.25
sd=1.5
Sample size
Power
Measure
dCor
MICe
Pearson
qad
RCD
rdc
Spearman
xicor

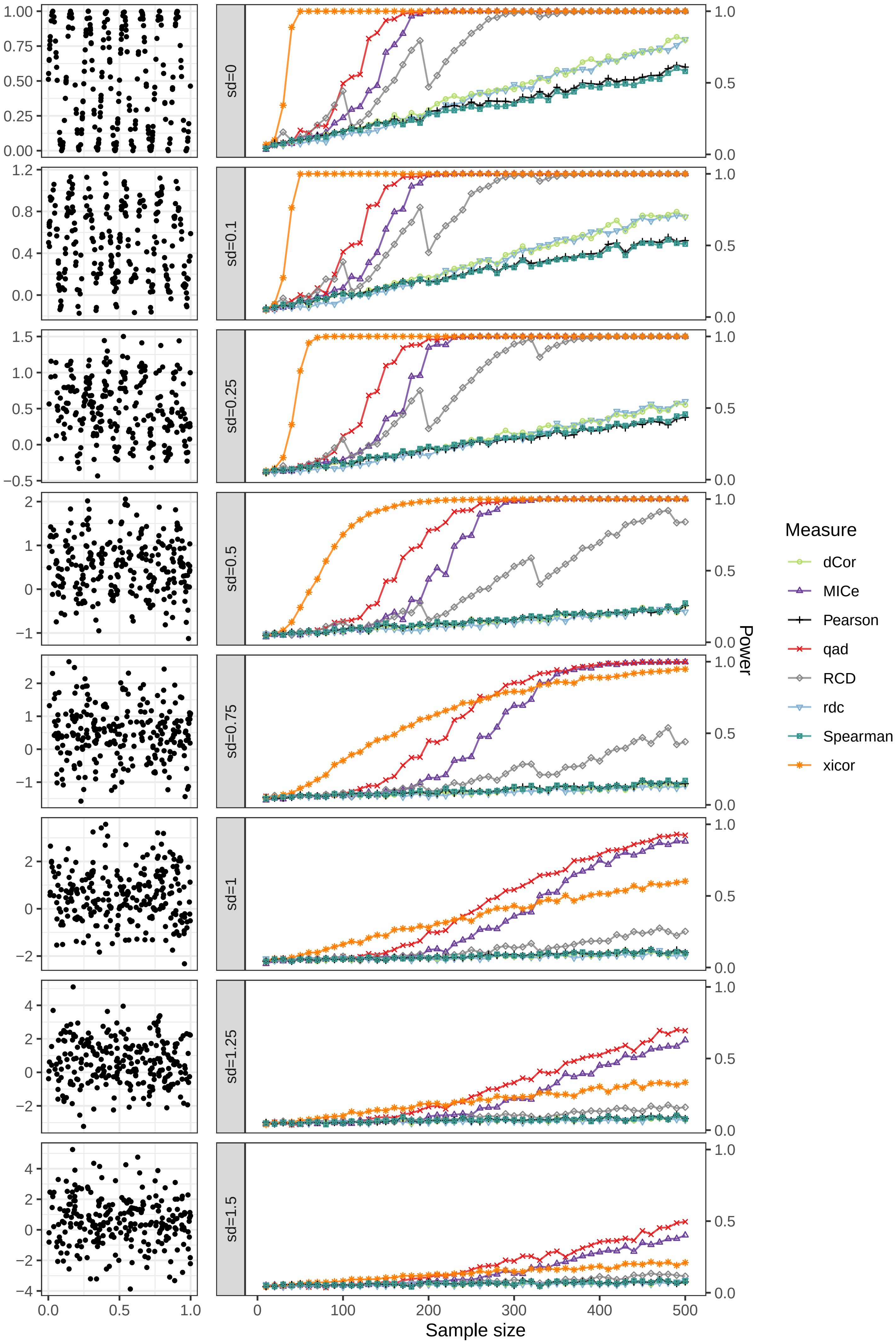
sd=0
sd=0.1
sd=0.25
sd=0.5
sd=0.75
sd=1
sd=1.25
sd=1.5
Sample size
Power
Measure
dCor
MICe
Pearson
qad
RCD
rdc
Spearman
xicor

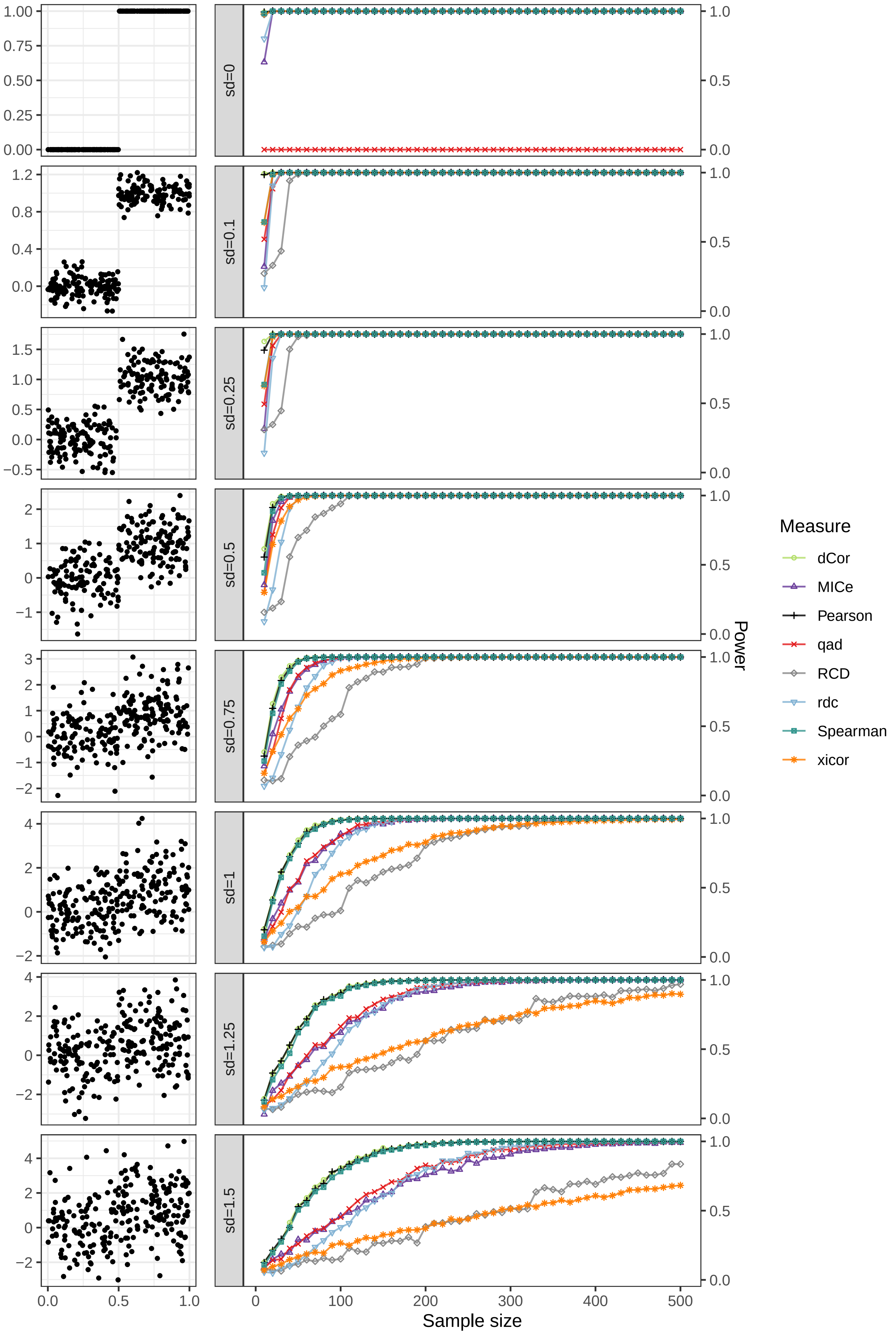

sd=0
sd=0.1
sd=0.25
sd=0.5
sd=0.75
sd=1
sd=1.25
sd=1.5
Power
Sample size
Measure
dCor
MICe
Pearson
qad
RCD
rdc
Spearman
xicor

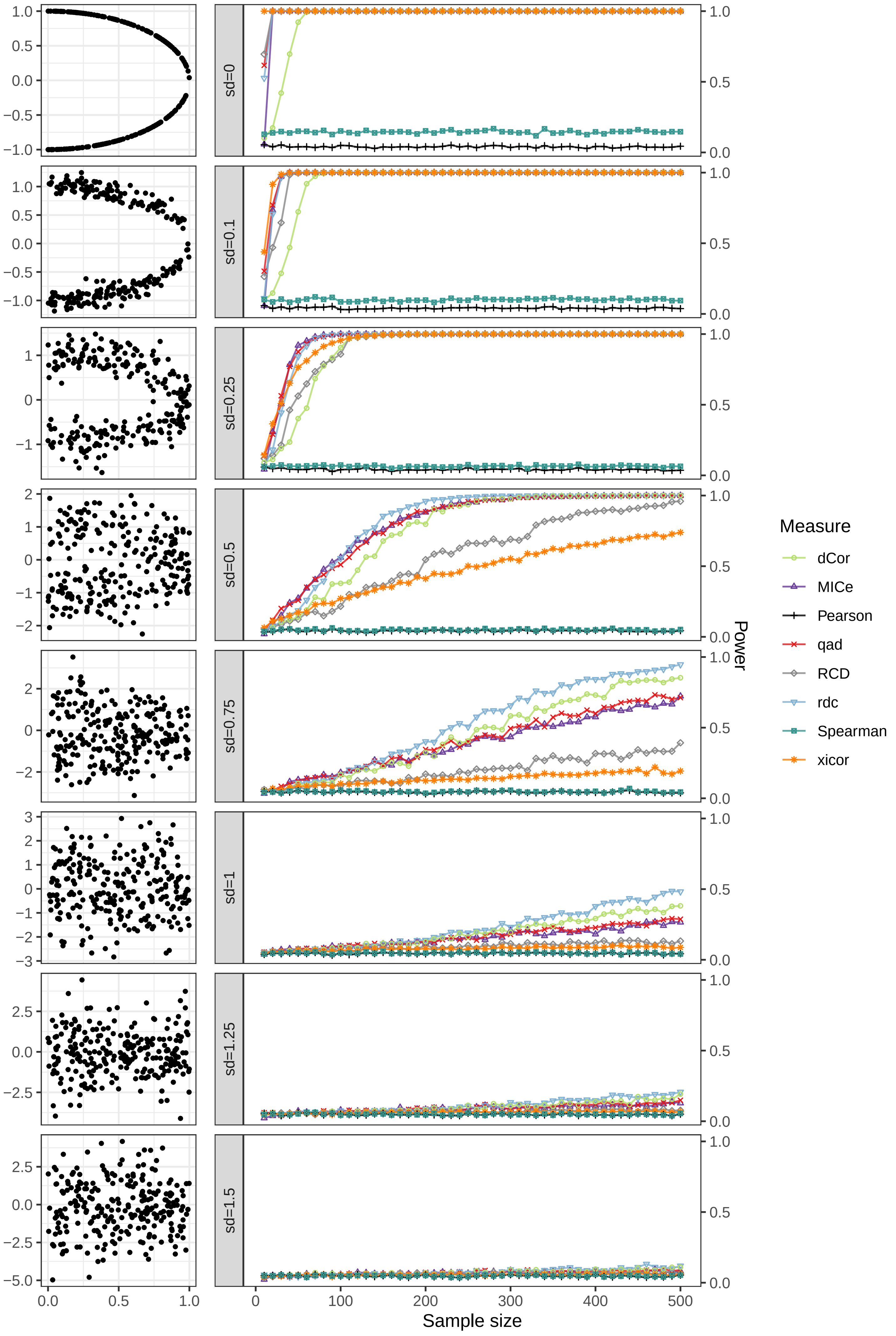

sd=0
sd=0.1
sd=0.25
sd=0.5
sd=0.75
sd=1
sd=1.25
sd=1.5
Sample size
Power
Measure
dCor
MICe
Pearson
qad
RCD
rdc
Spearman
xicor

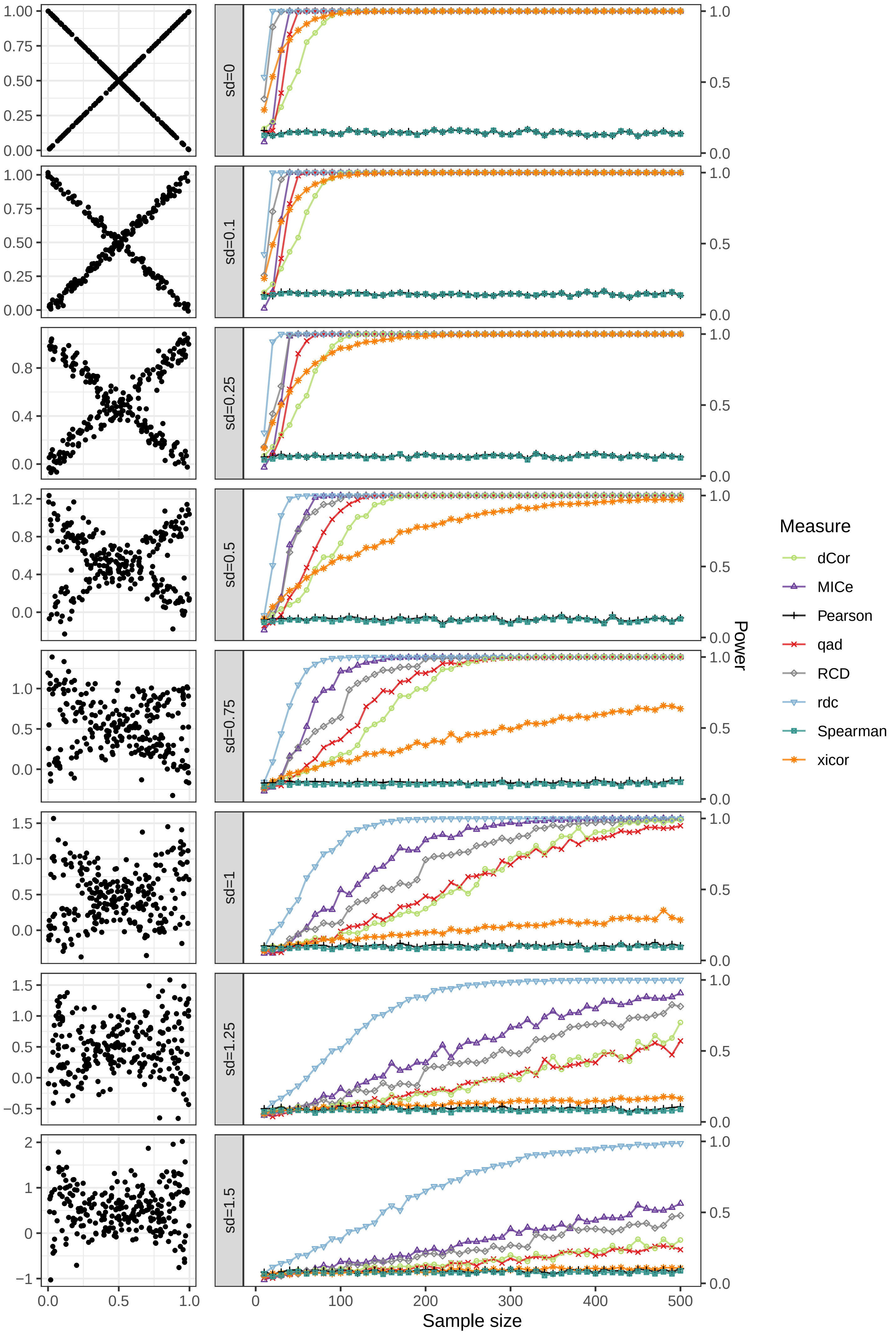

sd=0
sd=0.1
sd=0.25
sd=0.5
sd=0.75
sd=1
sd=1.25
sd=1.5
Sample size
Power
Measure
dCor
MICe
Pearson
qad
RCD
rdc
Spearman
xicor
