## SupplementaryInformation for "*qad:* An R-package to detect asymmetric and directed dependence in bivariate samples": Supplementary Information 1.R

```
#- [ Description ] ------------------------------------------------------------- 
#
# qad: An R-package to detect asymmetric and directed dependence in bivariate samples
#
# Florian Griessenberger (1), Wolfgang Trutschnig (1), Robert R. Junker (2,3,*)
#
# (1) Department of Mathematics, University of Salzburg, Salzburg, Austria
# (2) Department of Biosciences, University of Salzburg, Salzburg, Austria
# (3) Evolutionary Ecology of Plants, Faculty of Biology, Philipps-University Marburg, Marburg, Germany
# (*) Corresponding author


# Creation Date : 2021-02-03
# Source System : R version 4.0.2 (2020-06-22)
#---------------#
# Purpose of the script: Functions to generate the dependence structures
#_______________________________________________________________________________


#------------------------------------------------------------------------------#
#-------------------------- Extreme settings------------------------------------
#------------------------------------------------------------------------------#
#Noise was generated by drawing random numbers from a uniformly distributed random variable on the interval [-a,a], 
# whereby a was increased from 0 to 1 by steps of 1/100. We repeated the experiment R=1.000 times. 


#Dependence structures with samples in [0,1]^2 for the extreme settings
linear <- function(n, error = runif(n,-a,a)){
  x <- runif(n)
  y <- x + error
  A <- data.frame(x=x,y=y)
  A <- data.frame(x=(A$x-min(A$x))/(max(A$x)-min(A$x)) ,
                  y=(A$y-min(A$y))/(max(A$y)-min(A$y)))
  return(A)
}
quadratic <- function(n, error = runif(n,-a,a)){
  x <- runif(n, 0, 1)
  y <- 4*(x-0.5)^2 + error
  A <- data.frame(x=x,y=y)
  A <- data.frame(x=(A$x-min(A$x))/(max(A$x)-min(A$x)) ,
                  y=(A$y-min(A$y))/(max(A$y)-min(A$y)))
  return(A)
}
random <- function(n, error = runif(n,-a,a)){
  x <- runif(n)
  y <- runif(n) + error
  A <- data.frame(x=x,y=y)
  A <- data.frame(x=(A$x-min(A$x))/(max(A$x)-min(A$x)) ,
                  y=(A$y-min(A$y))/(max(A$y)-min(A$y)))
  return(A)
}
cubic <- function(n, error = runif(n,-a,a)){
  x <- runif(n, 0, 1)
  y <- 9*(x-1/3)^3 - 3*(x-1/3)^2- 1*(x-1/3) + 1/3 + error
  A <- data.frame(x=x,y=y)
  A <- data.frame(x=(A$x-min(A$x))/(max(A$x)-min(A$x)) ,
                  y=(A$y-min(A$y))/(max(A$y)-min(A$y)))
  return(A)
}
sinusoid <- function(n, error = runif(n,-a,a)){
  x <- runif(n, 0, 1)
  y <- 1/2*sin(4*x*pi) + 1/2 + error
  A <- data.frame(x=x,y=y)
  A <- data.frame(x=(A$x-min(A$x))/(max(A$x)-min(A$x)) ,
                  y=(A$y-min(A$y))/(max(A$y)-min(A$y)))
  return(A)
}
modulo1 <- function(n, error = runif(n,-a,a)){
  x <- runif(n, 0, 1)
  y <- (2*x) %% 1 + error
  A <- data.frame(x=x,y=y)
  A <- data.frame(x=(A$x-min(A$x))/(max(A$x)-min(A$x)) ,
                  y=(A$y-min(A$y))/(max(A$y)-min(A$y)))
  return(A)
}
torus <- function(n, error = runif(n,-a,a)){
  r <- sqrt(1) + error
  phi <- runif(n, 0, 2*pi)
  A <- data.frame(x=r*cos(phi),y=r*sin(phi))
  A <- data.frame(x=(A$x-min(A$x))/(max(A$x)-min(A$x)) ,
                  y=(A$y-min(A$y))/(max(A$y)-min(A$y)))
  return(A)
}
cross <- function(n, error = runif(n,-a,a)){
  x <- runif(n, 0, 1)
  y <- ifelse(rbinom(n, 1, prob = 0.5) == 0, x, 1-x) + error
  A <- data.frame(x=x,y=y)
  A <- data.frame(x=(A$x-min(A$x))/(max(A$x)-min(A$x)) ,
                  y=(A$y-min(A$y))/(max(A$y)-min(A$y)))
  return(A)
}
logistic<- function(n, error = runif(n,-a,a)){
  x <- runif(n, 0, 1)
  y <- 1/(1+ exp(-50*(x-0.5))) + error
  A <- data.frame(x=x,y=y)
  A <- data.frame(x=(A$x-min(A$x))/(max(A$x)-min(A$x)) ,
                  y=(A$y-min(A$y))/(max(A$y)-min(A$y)))
  return(A)
}


#------------------------------------------------------------------------------#
#----------------------Statistical power Analysis - SI -------------------------
#------------------------------------------------------------------------------#
#Dependence structures for the power analysis
generate_DS_sample <- function(n, 
                               type = "linear", 
                               noise_y = 0, 
                               x = runif(n, 0 , 1), 
                               noise_x = 0){
  #n....sample size 
  #type...type of dependence structure
  #noise...function of random noise
  
  #Generate x variable uniform within the interval [0,1] (here always without noise)
  x <- x + noise_x
  
  if(type == "random"){
    y <- runif(n,0,1)
  }
  if(type == "linear"){
    y <- x + noise_y
  } 
  if(type == "quadratic"){
    y <- 4*(x-0.5)^2 + noise_y
  }
  if(type == "cubic"){
    y <- 9*(x-1/3)^3 - 3*(x-1/3)^2- 1*(x-1/3) + 1/3  + noise_y
  }
  if(type == "sinus1"){
    y <- 1/2*sin(4*x*pi) + 1/2 + noise_y
  }
  if(type == "sinus2"){
    y <- 0.5*sin(16*x*pi) + 1/2 + noise_y
  }
  if(type == "doublesinus"){
    y <- 0.5*((2*rbinom(n,1,0.5)-1) * sin(3*pi*x)) + 0.5 + 1/4*noise_y
  }
  if(type == "4th-root"){
    y <- x^(1/4) + noise_y
  }
  if(type == "circle"){
    y <- 0.5*(2*rbinom(n, 1, prob = 0.5)-1) * (sqrt(1 - (2*x-1)^2)) + 0.5 + 1/4*noise_y
  }
  if(type == "U90degree"){
    y <- (2*rbinom(n, 1, prob = 0.5)-1) * (sqrt(1 - (x)^2)) + noise_y
  }
  if(type == "X"){
    y <- ifelse(rbinom(n, 1, prob = 0.5) == 0, x, 1-x) + 1/4*noise_y
  }
  if(type == "dnorm"){
    y <- dnorm(x, 0.5, 0.1)/4 + noise_y
  }
  if(type == "2xmod1"){
    y <- (2*x) %% 1 + noise_y
  }
  if(type == "5xmod1"){
    y <- (5*x) %% 1 + noise_y
  }
  if(type == "2boxes"){
    y <- ifelse(x < 0.5, 
                runif(length(x < 0.5), 0, 0.5), 
                runif(length(x >= 0.5), 0.5, 1)) + noise_y
  }
  if(type == "TwoLines"){
    y <- ifelse(rbinom(n, 1, prob = 0.5) == 0, x, 0.2*x) + noise_y
  }
  if(type == "logistic"){
    y <- 1/(1+ exp(-50*(x-0.5))) + noise_y
  }
  if(type == "step"){
    while(all(x<1/2) | all(x>=1/2)) x <- runif(n, 0, 1) + noise_x
    y <- ifelse(x< 1/2 ,0,1) + noise_y
  }
  
  return(data.frame(x,y))
}

noise <- function(n, sd = 1){
  #n... sample size
  #sd... standard deviation of normal distribution
  noise <- rnorm(n, mean = 0, sd = sd)
  return(noise)
}


#Example: 
df <- generate_DS_sample(500, type = "quadratic", noise_y = noise(500,0.05))
plot(df)
```
